# Structural basis for antagonism of the ubiquitin ligase BIRC6 by SMAC

**DOI:** 10.1101/2022.08.30.505748

**Authors:** Larissa Dietz, Cara J. Ellison, Carlos Riechmann, C. Keith Cassidy, F. Daniel Felfoldi, Adán Pinto-Fernández, Benedikt M. Kessler, Paul R. Elliott

## Abstract

Apoptosis, a form of genetically programmed cell death, can be triggered by either internal or external signals ultimately activating caspases, a family of proteases^1^. Certain members of the inhibitors of apoptosis (IAP) family are sentinel proteins preventing untimely cell death by inhibiting caspases. IAPs are in turn regulated by antagonists including second mitochondria-derived activator of caspase (SMAC). Baculoviral IAP repeat-containing protein 6 (BIRC6), a giant IAP, possesses dual E2/E3 ubiquitin ligase activity and is implicated in apoptosis via caspase inhibition^2–7^. How this is achieved remains unknown. Here we show BIRC6 directly restricts activated caspase-3, and ubiquitinates activated caspases-3, −7 and −9 working exclusively with the non-canonical E1, UBA6. Importantly, we show SMAC supresses both mechanisms. Cryo-electron microscopy (cryo-EM) structures of BIRC6 alone and in complex with SMAC reveal BIRC6 exists as an anti-parallel dimer with a substrate-binding module juxtaposed to the catalytic domain at each end, and we identify multiple highly conserved unannotated domains important for architecture and function. Through our structural, biochemical and biophysical findings, we discover SMAC engages BIRC6 at multiple sites resulting in a sub-nanomolar affinity enabling SMAC to competitively displace caspases, thus antagonising BIRC6-mediated caspase inhibition.

---

Programmed cell death is required for normal development but must be tightly controlled. Several mechanisms exist to prevent cell death in the absence of signalling cues and ensure caspase activity is stringently regulated. Firstly, caspases are expressed as inactive zymogens (procaspases) requiring proteolytic processing for activation, and secondly, a subset of IAPs restrict caspase activity.

IAPs contain a signature baculoviral IAP repeat (BIR) domain that bind caspases amongst other substrates and, frequently, a C-terminal ubiquitin ligase domain responsible for attaching ubiquitin post-translationally to target proteins. Of the eight mammalian IAPs, only three are well-characterised inhibitors of apoptosis – cellular IAP1 (cIAP1, BIRC2), cIAP2 (BIRC3) and X chromosome-linked IAP (XIAP, BIRC4)^8^. XIAP directly inhibits activated caspases-3, −7 and −9 via a conserved Asp-containing pocket in BIR2 and BIR3 domains binding to the amino terminus of the processed caspase small subunit^9^. cIAP1/2 do not physically restrict caspase activity but promote caspase degradation through ubiquitination, and promote cell survival through production of pro-inflammatory ubiquitin chains in inflammatory signalling pathways^10–12^.

BIRC6 (BRUCE, Apollon) is a giant IAP (4,857 amino acids) conserved from Drosophila to humans containing the hallmark BIR domain and a ubiquitin conjugation domain (UBC) conferring E2 ubiquitin ligase activity. Genetic studies highlight an essential role of BIRC6 in development and apoptosis inhibition – *BRUCE* knock-out mice are embryonically lethal owing to placental defects or increased apoptosis levels^7, 13^. Further, cellular studies detected dual E2 and E3 ubiquitin ligase activity (only identified in one other ligase^14^) and inhibition of caspases −3, −7 and −9^2, 4^. Roles of BIRC6 in other cellular processes such as autophagy are also emerging^15–18^.

IAPs themselves must be inhibited to drive apoptosis, achieved in the intrinsic pathway by two mitochondrial-released proteins: HtrA2 cleaves XIAP and cIAP1/2^19, 20^, whilst SMAC directly competes with caspases binding to XIAP^21, 22^. BIRC6-mediated inhibition of apoptosis has been shown to be suppressed by SMAC in cellular studies^2^ through uncharacterised mechanisms.

Despite this clear cellular and genetic evidence for BIRC6 playing a critical role in inhibiting apoptosis, understanding of BIRC6 ubiquitin ligase activity, anti-apoptotic mechanisms, and antagonism by SMAC remain unknown.

## BIRC6 functions exclusively with UBA6

To identify BIRC6 regulatory proteins, we expressed and purified recombinant full-length BIRC6 (**Extended Data Fig. 1a**) and performed mass spectrometry on affinity-purified complexes from unstimulated HEK293F cells. This approach revealed significant enrichment of the non-canonical E1, UBA6 (**Fig. 1a**). Two E1 ubiquitin ligases exist with UBA1 responsible for initiating the majority of ubiquitination cascades. UBA6, however, functions with a small subset of E2 enzymes and facilitates the transfer of the ubiquitin-like protein FAT10 in addition to ubiquitin^23, 24^. Strikingly, BIRC6 only receives ubiquitin from UBA6 and not UBA1 in in vitro transthiolation reactions (**Fig. 1b, Extended Data Fig. 1b**). BIRC6 weakly accepts FAT10 from UBA6, compared to ubiquitin, and to a much lesser extent compared to UBE2Z, the known FAT10 acceptor^25^ (**Extended Data Fig. 1c**). Our findings place BIRC6 as the only UBA6-specific, ubiquitin-selective ligase (**Extended Data Fig. 1d**).

**Fig. 1.**
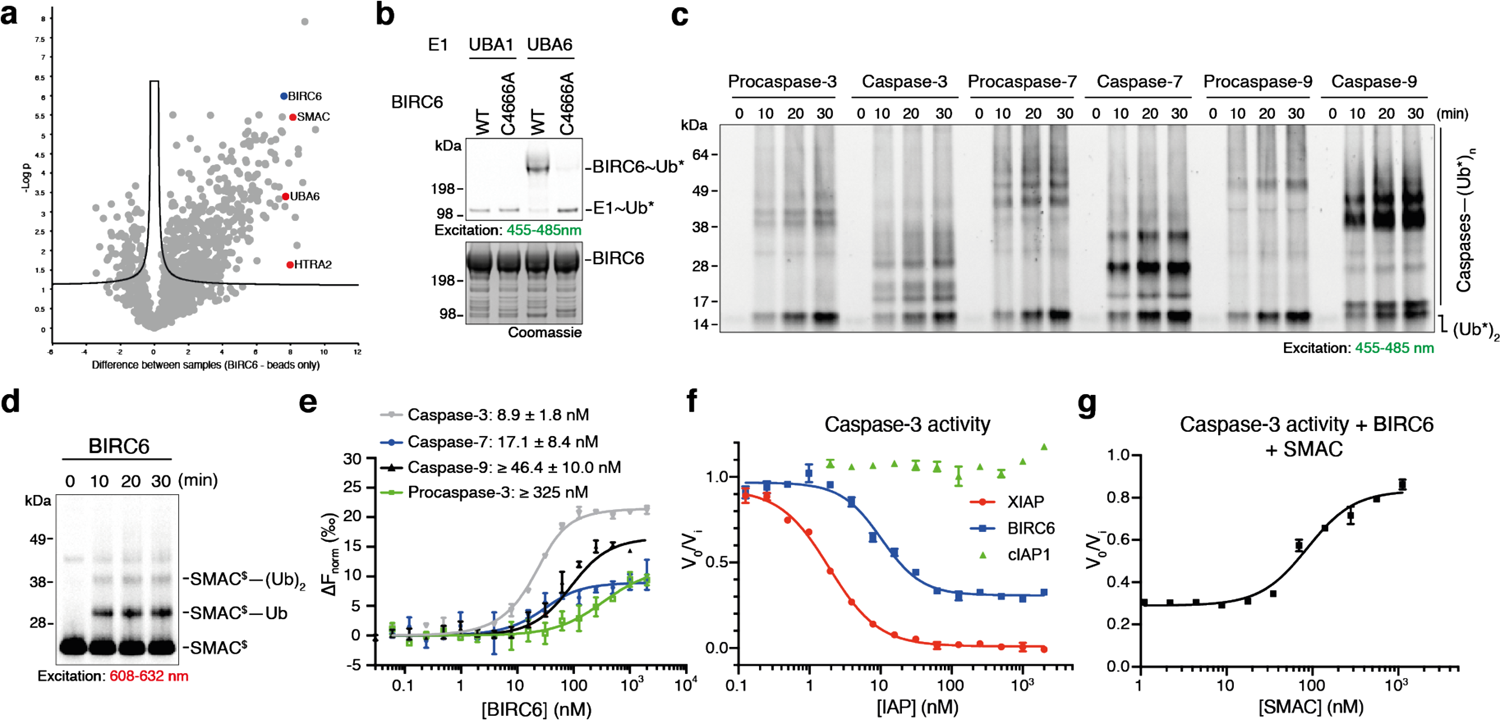
BIRC6 is a regulator of apoptosis by direct and indirect inhibition of caspases. **a,** Identification of BIRC6 interaction partners using affinity-purified MS/MS. Volcano plot depicts significant enrichment of proteins bound to BIRC6 compared to a beads-only control. **b**, BIRC6 receives activated ubiquitin from UBA6 and not UBA1 in an in vitro transthiolation reaction analysed on a non-reducing SDS-PAGE gel. WT, wild-type; C4666A, catalytically inactive mutant; ∼ denotes thioester bond. Representative of 3 independent repeats. **c**, In vitro ubiquitination by BIRC6 of initiator and executioner caspases visualised using BDP-labelled ubiquitin (*). Gel representative of 3 independent repeats. **d**, BIRC6 ubiquitinates a key regulator of intrinsic apoptosis, SMAC, shown through an in vitro ubiquitination assay as in **c**, but using Cy5-labelled SMAC (^$^); – denotes covalent bond. Representative of at least 3 independent repeats. **e**, BIRC6 binds active caspases with high affinity, measured by microscale thermophoresis (MST). Data represents 2 independent replicates. **f**, BIRC6 directly inhibits caspase-3 activity in a fluorogenic substrate cleavage assay. Graphs show rate of substrate cleavage by active caspase-3 in the presence of IAPs, normalised to caspase-3 in the absence of IAPs. Results represent 2 independent repeats. **g**, SMAC alleviates BIRC6-mediated caspase-3 inhibition in a fluorogenic substrate cleavage assay recording caspase-3 activity in the presence of 50-fold molar excess BIRC6 to caspase-3 with varying SMAC concentrations normalised to caspase-3 activity alone. Results represent 2 independent repeats. Error bars represent standard deviation (SD).

Next we asked whether BIRC6 E2 activity is restricted to specific E3 families. In in vitro auto-ubiquitination assays testing a panel of E3s representative of the three families, BIRC6 displays cross-family E2 activity working with UBE3C and ARIH1 of the HECT and RBR E3 families, respectively (**Extended Data Fig. 1e-g**). With over 600 known E3 ubiquitin ligases, it is likely that BIRC6 also functions as a stand-alone E2 to additional E3 ubiquitin ligases.

## BIRC6 restricts caspase activity

We also identified strong enrichment of the intrinsic apoptosis antagonists SMAC and HtrA2 in complex with BIRC6 (**Fig. 1a**) leading us to test BIRC6 E2/E3 activity in ubiquitinating critical components of intrinsic apoptosis. We observed robust ubiquitination of the activated intrinsic initiator caspase, caspase-9, and the activated executioner caspases −3 and −7 with weak ubiquitination of procaspases (**Fig. 1c, Extended Data Fig. 2a**). Additionally, BIRC6 ubiquitinates SMAC and inactive HtrA2 (S306A) (**Fig. 1d, Extended Data Fig. 2b**) but does not ubiquitinate activated caspase-8, the initiator caspase of extrinsic apoptosis (**Extended Data Fig. 2c**). In all instances, BIRC6 catalyses multi-monoubiquitination of substrates, in contrast to polyubiquitination by XIAP and cIAP1 (**Extended Data Figs. 2d-g**). Despite FAT10 loading onto BIRC6, we did not detect any transfer of FAT10 onto these substrates by BIRC6 (**Extended Data Fig. 1h**). Our results show BIRC6 functions as a combined E2/E3 ubiquitin ligase ubiquitinating key players in intrinsic apoptosis.

**Fig. 2.**
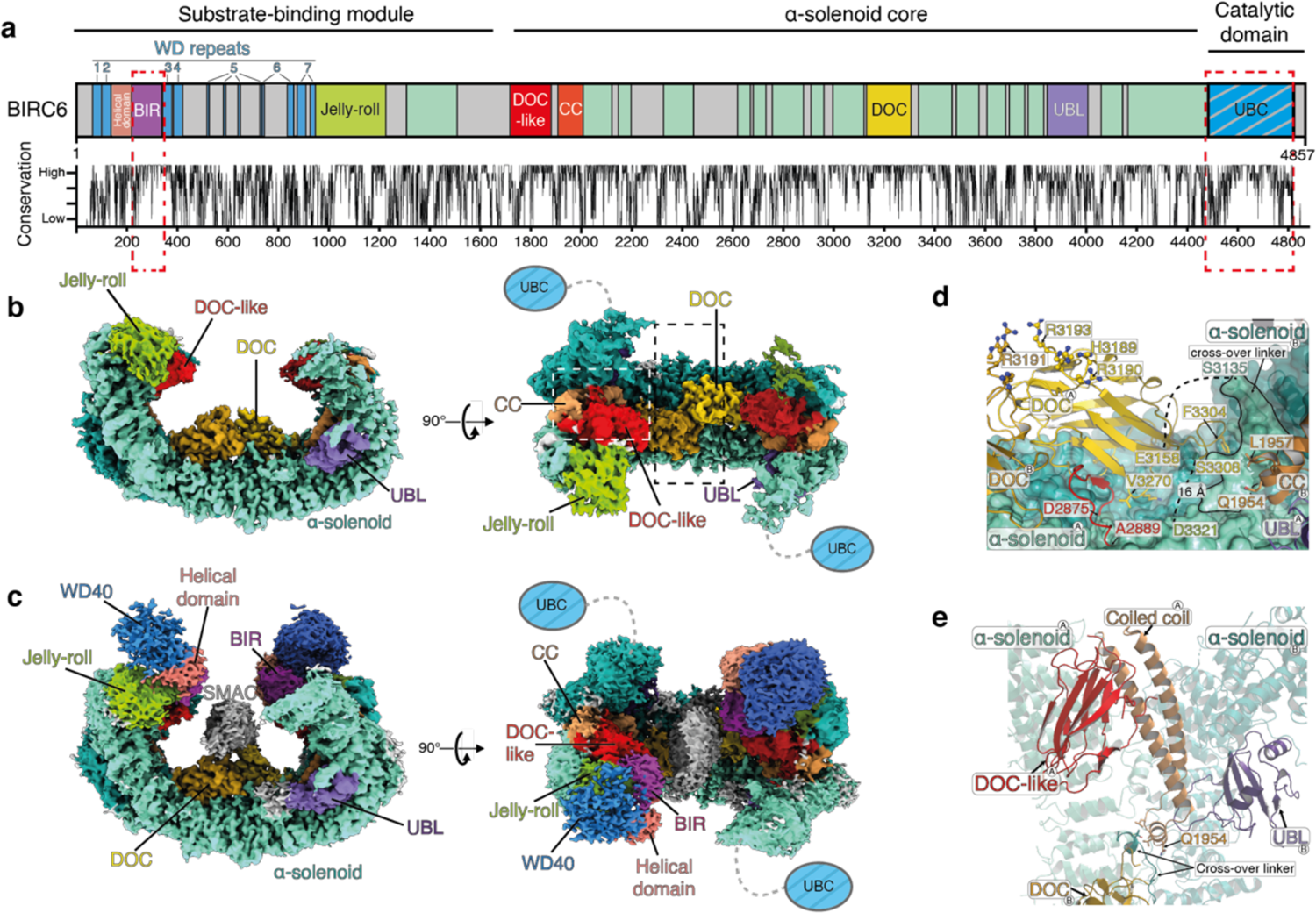
The anti-parallel dimeric architecture and intricate domain arrangement of BIRC6 is important for SMAC engagement. **a,** New domain annotation of BIRC6 revealed through our study. Corresponding residue conservation of BIRC6 amongst 126 orthologues is shown below. Previously known BIR and UBC domains, are highlighted in red dashed boxes. Areas in grey are not visible in our cryo-EM structures. b, Composite cryo-EM density representing BIRC6 contoured at 3σ. Domains coloured according to a. Black and white dashed boxes highlight enlarged regions in d, and e, respectively. c, BIRC6 in complex with SMAC shown by cryo-EM density contoured at 2σ. Molecular Dynamics Flexible Fitting was used to fit AlphaFold2^40^ models of SMAC, the WD40, BIR and helical domains into the low threshold regions of the map. Domains coloured according to a. d, Magnification of the central DOC domain residues Ile3160 – Ser3308 (yellow) showing the unique protruding loop and inter-chain cross-over from a side view with the UBL (purple) and coiled-coil, CC, (bronze) domains to the right. Chains to which the domains belong are indicated by ^A^ and ^B^. Red denotes residues Asp2875 – Ala2889 rising from the α-solenoid core to stabilise the DOC domain positioned over the adjacent chain. Dashed lines indicate flexible loops between DOC and α-solenoid domains, the loop between Ser3308 and Asp3321 bridges 16 Å. e, Coiled coil (bronze) domain stabilises the BIRC6 dimer by contacting the α-solenoid core and the DOC-like domain (red) of the same chain in addition to the UBL (purple) domain of the opposite chain. Corresponding chains are indicated as in d. Additional contacts are made between the hinge of the coiled coil (Gln1954) to the cross-over linker for the DOC domain, as shown in d.

As we observed robust ubiquitination of activated caspases by BIRC6, we tested whether BIRC6 can directly inhibit caspase activity. We recorded tight, nanomolar, association between BIRC6 and activated caspases-9, −3 and −7 (Kd 46.4 ± 10 nM, 8.9 ± 1.8 nM, 17.1 ± 8.4 nM, respectively) (**Fig. 1e**). Addition of stoichiometric excess of BIRC6 impaired activated caspase-9 activity but only in the absence of the apoptosome complex (**Extended Data Fig. 2h-k**), the macromolecular complex responsible for caspase-9 activation^26^. In contrast, we observed BIRC6 directly inhibits caspase-3 activity in a fluorogenic cleavage assay (**Fig. 1f**) and weakly inhibits caspase-7 activity (**Extended Data Fig. 2l**). In summary, our data reveal dual anti-caspase activity of BIRC6: BIRC6 directly inhibits activated caspase-3, and indirectly inhibits activated caspases by multi-monoubiquitination, leading to caspase degradation in a cellular context.

## BIRC6 architecture defines antagonism by SMAC

We next explored how SMAC regulates the anti-apoptotic activity of BIRC6. In a competition fluorogenic cleavage assay, we find that BIRC6-mediated caspase-3 inhibition reduces as the concentration of SMAC increases (**Fig. 1g**), showing SMAC impairs BIRC6-caspase-3 inhibition. To reveal how this antagonism is achieved, we used single-particle cryo-EM analysis to determine the structures of BIRC6 alone or in complex with SMAC (**Fig. 2a-c, Extended Data Figs. 3, 4, Supplementary Data Table 1**). Our structures reveal that BIRC6 possesses an overall crab-like architecture comprising an anti-parallel dimer, consistent with SEC-MALS (**Extended Data Fig. 3e**), consisting of a rigid α-solenoid core (residues 1010-4502) (resolved to 2.9 Å) (**Extended Data Fig. 6a,b**) with a flexible N-terminal substrate-binding module and a highly flexible C-terminal catalytic domain acting as antennae and claws, respectively, guiding potential substrates towards a central mouth-like cavity (**Fig. 2b,c**). The anti-parallel arrangement juxtaposes the substrate-binding module with the catalytic ubiquitin ligase domain at each end, potentially allowing for both intra- and inter-chain activity.

**Fig. 3.**
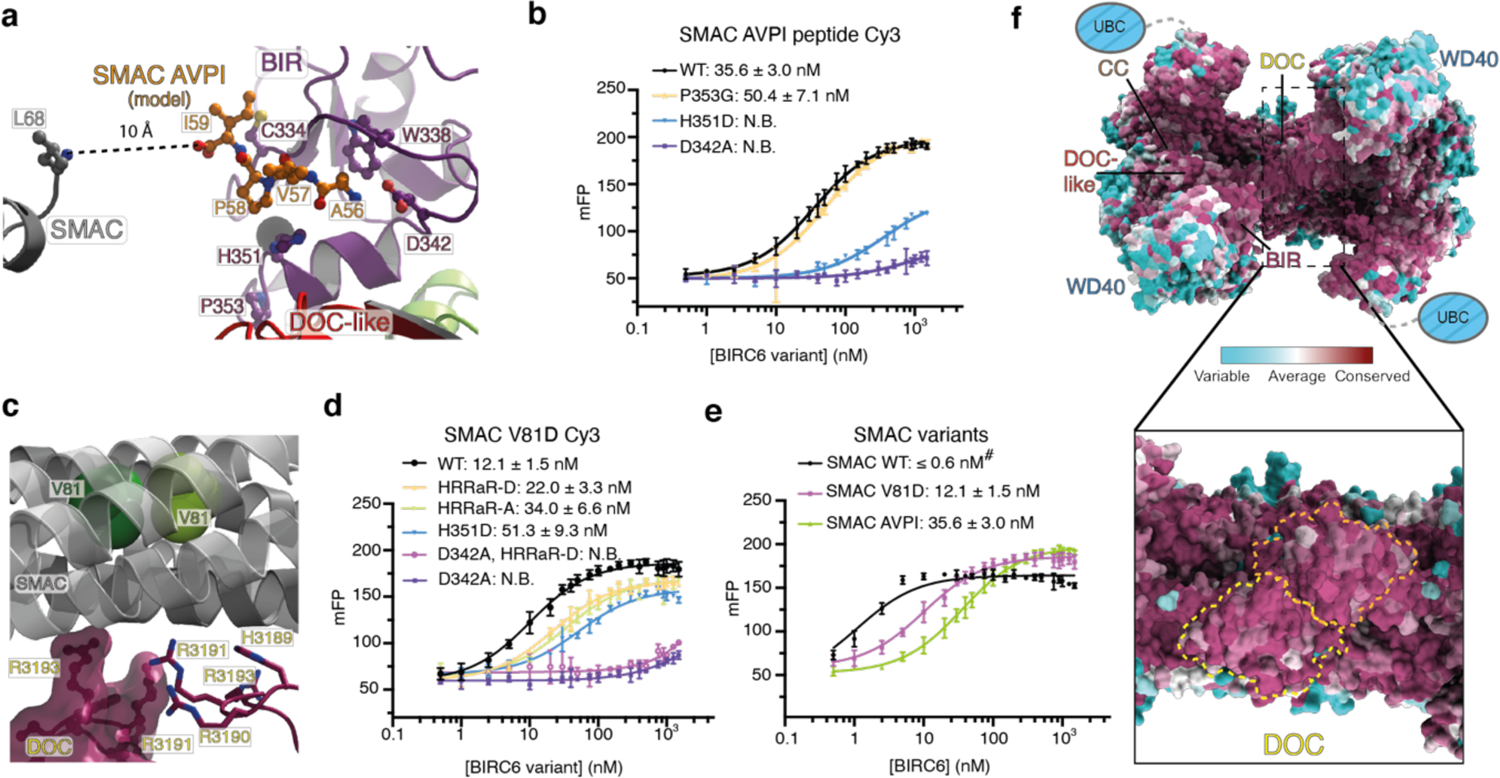
Dimeric SMAC binds to BIRC6 via a multi-site mechanism. **a,** Position of SMAC N-terminal peptide (orange) binding to BIRC6 BIR domain (purple) modelled by Molecular Dynamics Modelling. The peptide C terminus is 10 Å from the N terminus of the SMAC helical bundle (Leu68) (grey), providing sufficient distance for seven amino acids to span. **b**, Affinities of BIRC6 wild-type (WT) and variants to Cy3-labelled SMAC N-terminal peptide determined using fluorescence polarisation (FP). Data represents two independent repeats in triplicate. **c**, The DOC domain of BIRC6 provides an additional interface for SMAC binding involving a highly conserved positively charged sequence at the tip of the DOC domain (residues 3189-3193;HRRaR). **d**, Affinity measurements between Alexa 555 monomeric SMAC (V81D) and BIRC6 variants determined by FP. Results are representative of at least 2 independent repeats. **e**, Determination of BIRC6 affinity to labelled SMAC (WT), SMAC monomer (V81D) and SMAC peptide by FP. # denotes the BIRC6-SMAC (WT) affinity being tighter than the minimum concentration of labelled SMAC detectable. Results are representative of 3 independent repeats in triplicate. Error bars represent SD. **f**, Conservation analysis of 126 BIRC6 orthologues with a magnification of the highly conserved central DOC domain outlined in yellow. Highly conserved residues are shown in maroon, variable residues in cyan, as defined by ConSurf^41^.

**Fig. 4.**
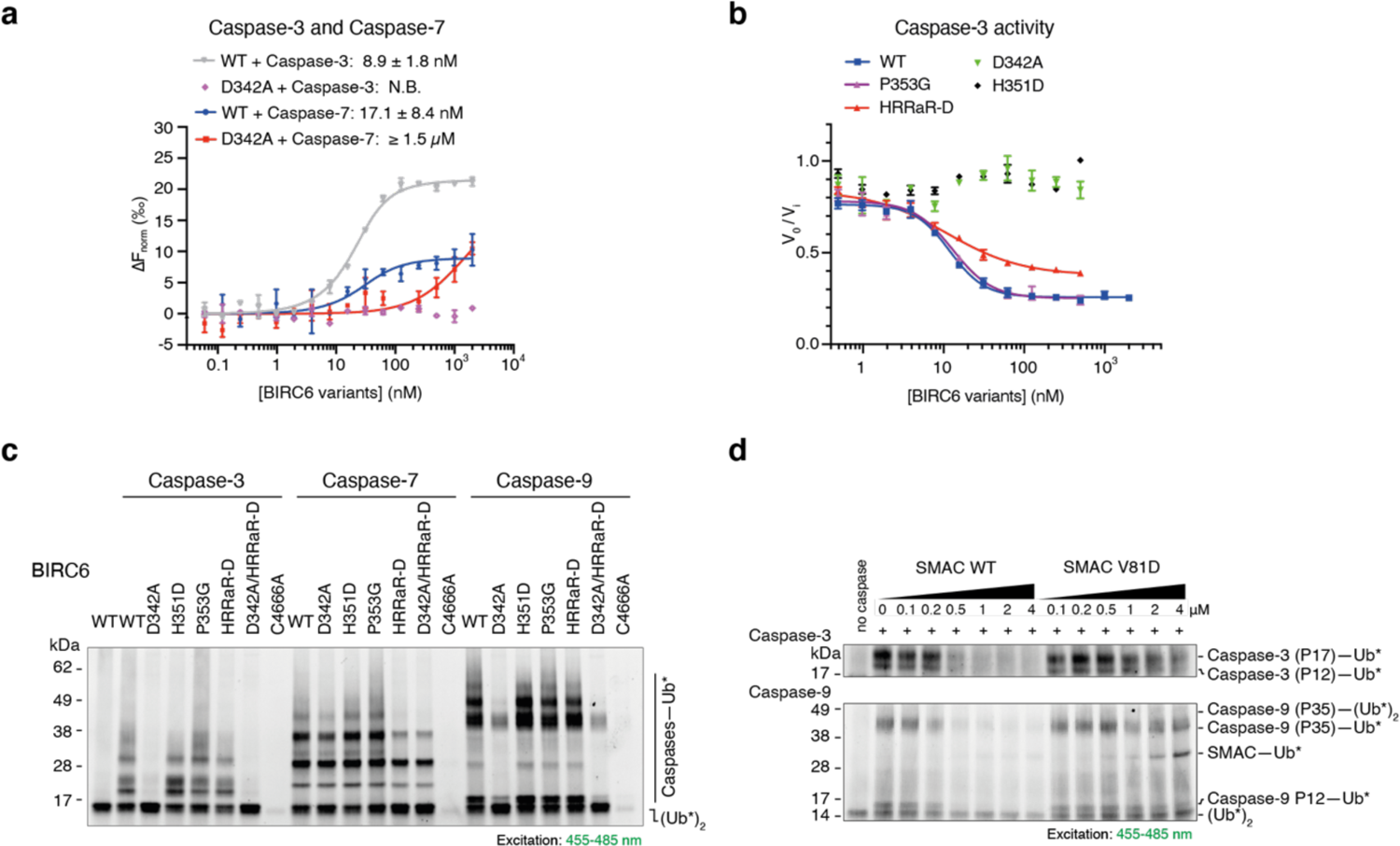
Caspases and SMAC bind to BIRC6 through overlapping sites. **a**, Affinities between BIRC6 WT or mutant D342A and RED-tris-NTA labelled caspase-3 and −7 measured using microscale thermophoresis (MST). Results are representative of 2 independent repeats. **b**, Mutations in the BIRC6 BIR domain SMAC peptide-binding pocket abolish direct inhibition of caspase-3 activity in a fluorogenic substrate cleavage assay. Plots indicate rate of substrate cleavage by active caspase-3 in presence of BIRC6 variants normalised against caspase-3 activity alone. Error bars represent SD. **c**, In vitro ubiquitination assays using BDP-labelled ubiquitin (*) comparing the effect of BIR domain and DOC domain mutations on BIRC6 ubiquitination of activated caspases. Gel is representative of 3 independent repeats. **d**, SMAC dimerization enables inhibition of BIRC6 ubiquitination of activated caspase-3 and −9. Increasing concentrations of SMAC dimer (WT) or monomer (V81D) were incubated with BIRC6 prior to addition of activated caspase-3 or caspase-9 in an in vitro ubiquitination reaction using BDP-labelled ubiquitin (*). Gels are representative of 2 independent repeats.

Our structures reveal multiple, previously unannotated domains interwoven around the solenoid core, contributing to BIRC6 architecture and function (**Fig. 2a**). In the central cavity, we identify a DOC domain (residues 3160-3308), an all-beta strand structure first identified for substrate-binding in the APC/C APC10 subunit^27^, and since identified in other E3 ubiquitin ligases^28^ (**Extended Data Fig 5a-c**). We observe a structural role for the central DOC domain; a short loop from the core of one polypeptide chain snakes over and nestles within the solenoid core of the adjacent chain, positioning the DOC domain on top of this adjacent chain, inter-locking the two. The DOC domain is further stabilised by a beta strand (residues 2875-2889) protruding ∼250 residues prior to the DOC domain (**Fig. 2d**). In the BIRC6-SMAC complex, we observed density (∼8 Å) above this central DOC domain allowing us to place the dimeric structure of SMAC in this pivotal position (**Fig. 2c**). In contrast to other DOC domain proteins, BIRC6 possesses a unique loop extending from the DOC domain tip, which makes contacts with the SMAC helical bundle (**Fig. 3c, Extended Data Fig. 5c**), indicating this unique extension is important for substrate engagement.

In the N-terminal arm, we identify a DOC-like domain and a jelly-roll domain both with potential for substrate-binding and positioning in the central cavity. Towards the C-terminal arm, we identify a UBL domain, containing a long unstructured loop insertion (**Extended Data Fig. 6c,d**). Of central importance to the structural arrangement of these domains is an anti-parallel coiled-coil running the length of the arm, tucked between the anti-parallel solenoid core and making contacts with the DOC-like and UBL domains, the latter from the adjacent chain (**Fig. 2e, Extended Data Fig. 6e,f**). In addition, the elbow of the coiled coil (Gln1954 and Leu1957) forms contacts to the incoming helix leading to the DOC domain. Together, the coiled coil thus acts as a structural hinge stabilising the intricate interwoven domain arrangement.

Our SMAC-bound BIRC6 structure revealed further domains in the N-terminal substrate-binding module including a WD40 domain formed over 1,000 residues interrupted by a short helical domain and BIR domain (**Extended Data Fig. 7a-d**). This interrupted WD40/BIR domain assembly forms a tightly packed structural element and further expands its repertoire of potential substrate interaction regions (**Fig.2a**).

We extended our analysis to other BIRC6 orthologues revealing all these structural features are conserved (**Extended Data Figs. 8, 9**), supporting the conclusions that the overall architecture and domain composition is essential for BIRC6 function.

## SMAC binding to BIRC6

We examined the interactions between BIRC6 and SMAC in more detail. Our BIRC6-SMAC structure shows dimeric SMAC positioned above the DOC domain (**Fig. 2c, Extended Data Fig. 7a,e**). The start of the SMAC helical bundle (Leu68) points towards the BIR domain (**Fig. 3a**) indicating a potential interaction with the N terminus of mature SMAC. Through molecular docking simulations, SMAC N-terminal AVPI sequence binds to a well-defined pocket in BIRC6 BIR domain with an invariant residue Asp342 hydrogen bonding to SMAC N-terminal Ala56 (**Fig. 3a**). In support of this model, we recorded a strong, nanomolar interaction between BIRC6 and SMAC N-terminal peptide using fluorescence polarisation (FP) and mutation of Asp342 or His351 in full-length BIRC6 abrogated binding to SMAC peptide (**Fig. 3b, Extended Data Fig. 10a**); mutating Pro353 positioned on the external face of the peptide-binding pocket did not affect peptide-binding affinity (**Fig. 3b**). In conclusion, the N-terminal residues of SMAC bind to the BIR domain via the conserved Asp342-containing pocket.

BIRC6 binding to monomeric SMAC is three times stronger (Kd 12.1 ± 1.5 nM) than to the isolated SMAC peptide (Kd 35.6 ± 3.0 nM) (**Fig. 3d**) and is driven through the Asp342-containing pocket (loss of Asp342 abrogated binding (**Fig. 3d**)). Consistent with our complex structure, we detect a binding contribution from the DOC domain protruding loop as mutating the positively charged DOC domain tip (residues 3189-3193, HRRaR to all Ala or all Asp) reduced the binding affinity to that measured between BIRC6–SMAC N-terminal peptide (**Fig. 3c,d, Extended data Fig. 7f**).

Dimeric, WT, SMAC binds BIRC6 with a sub-nanomolar affinity estimated via a competition assay with SMAC peptide (**Fig. 3e, Extended data Fig. 10b**), in agreement with the SMAC-XIAP BIR2-BIR3 affinity reported^29^. Mutation of D342A in combination with HRRaR–D reduced, but did not abolish, binding to dimeric SMAC (**Extended data Fig. 10c**) suggesting SMAC dimerization strengthens additional weaker multi-valent interactions with BIRC6.

Together, our biophysical analyses support dimeric SMAC binding to dimeric BIRC6 is multi-site, driven through the BIR domain with additional contributions from the DOC domain and other interfaces. Conservation analyses amongst a large range of BIRC6 orthologues reveals a highly invariant inner face incorporating the DOC domain (**Fig. 3f, Extended Data Fig. 9**) indicating essentiality for BIRC6 substrate recognition.

## SMAC antagonises BIRC6 caspase-binding

We then asked what role the BIRC6-SMAC binding mechanism plays in the inhibition of caspases. First, we investigated whether BIRC6-caspase binding involves the same key domains and residues as SMAC binding. Indeed, BIR domain D342A or H351D mutations substantially reduced, or even abrogated, BIRC6 binding to activated caspases-3, −7 and −9 (**Fig. 4a, Extended Data Fig. 10d**). Importantly, BIRC6 D342A or H351D no longer inhibited caspase-3 activity and mutating DOC HRRaR–D also reduced caspase-3 inhibition (**Fig. 4b**). Additionally, BIRC6 D342A displayed reduced ubiquitination of caspases-3 and −9 (**Fig.4c**). Mutating Pro353, corresponding to Pro325 important for XIAP-caspase-9 binding^30^ but positioned away from the BIRC6 SMAC peptide-binding pocket (**Fig. 3a**), did not affect BIRC6-caspase-9 binding or ubiquitination (**Fig. 4c, Extended Data Fig. 10d**). In conclusion, not only are the BIR and DOC domains also involved in BIRC6 inhibition of caspases-3 and −9, but critically, residues pivotal for caspase binding are identical to those bound by SMAC, suggesting SMAC directly competes with these caspases for BIRC6-binding. Curiously, BIR domain mutations did not markedly reduce caspase-7 ubiquitination; however, mutating DOC HRRaR–D did reduce ubiquitination (**Fig. 4c**), suggesting a greater contribution from the DOC domain in binding activated caspase-7.

Finally, we assessed whether the affinity of SMAC binding to BIRC6 contributes to its inhibition. Dimeric SMAC binds BIRC6 with at least 10 times greater affinity than that of monomeric SMAC (V81D) (**Fig. 3e**). Strikingly, monomeric SMAC (V81D) showed reduced ability to antagonise BIRC6 ubiquitination of caspases-3 and −9 compared to the dimeric form, even at higher SMAC concentrations (**Fig. 4d, Extended Data Fig. 10e,f**). Therefore, the multi-site binding of dimeric SMAC to dimeric BIRC6 we observe in our cryo-EM structure results in a sub-nanomolar affinity key for the ability of SMAC to block BIRC6 anti-apoptotic function.

## Implications for BIRC6 function

These findings cement BIRC6 as a bona fide apoptosis inhibitor. BIRC6 physically restricts caspase-3 activity, positioning BIRC6 alongside XIAP as the only IAPs with this behaviour^30–33^. Furthermore, BIRC6 combines this inhibitory activity with that of cIAP1/2 through ubiquitinating activated caspases^4, 12, 34^; BIRC6 may collude with another E3 ubiquitin ligase extending the initially deposited monoubiquitin leading to caspase degradation in a cellular context (**Extended Data Fig. 10g,h**).

We uncover SMAC antagonises BIRC6 through directly out-competing caspases bound to key residues in conserved domains. SMAC dimerization and the constitutive anti-parallel dimeric nature of BIRC6 are critical for this through a resultant sub-nanomolar affinity complex. For the first time our structures reveal an additional antagonistic role of the SMAC central helical bundle in IAP inhibition. In addition to it enabling SMAC dimerization, key for simultaneous binding of BIR2 and BIR3 domains within XIAP^21, 22^ and cIAP1/2 and critical for engagement of the resultant two BIRC6 BIR domains revealed through our structure, we identify an occlusion role: interactions of the SMAC helical bundle with the BIRC6 DOC domain further increase the interaction affinity and physically obstruct other substrates from binding in the central cavity.

IAPs, including BIRC6, are frequently over-expressed in cancers^35–38^ and detailed understanding of XIAP and cIAP1/2 inhibitory activity and regulation has enabled design of therapeutics exploiting the ability of SMAC to block XIAP and cIAP1/2 function to restore apoptosis^39^. Our findings revealing the mechanisms of BIRC6 anti-apoptotic activity and multi-modal inhibition by SMAC now provide a biochemical and structural framework for future design of small molecule inhibitors targeting this IAP.

Finally, discovery of BIRC6 UBA6-dependent ubiquitination activity opens exploration of BIRC6 control of emerging UBA6-regulated pathways^17, 23^, potentiated by the array of highly conserved substrate-recruitment domains we identify. Its ability to function as a stand-alone E2 in addition to a dual E2/E3 further extends the reach of BIRC6 through co-operating with specific E3 ubiquitin ligases.

## Methods

### Protein expression and purification from Sf9 cells

Full-length BIRC6 codon-optimised for expression in *Spodoptera frugiperda* (Sf9) cells was assembled from six synthetic DNA fragments (Thermo Fisher) and cloned using Gibson Assembly (NEB) into a modified pACEBac1 vector containing a Twin-Strep tag®, mScarlet and a 3C protease site. Full-length UBA6 and Apaf1, codon-optimised for Sf9 cell expression, were cloned into this modified pACEBac1 vector. All plasmids were verified by Sanger sequencing. Vectors were transformed into DH10Bac cells (Invitrogen). Bacmids were isolated and transfected into Sf9 cells to produce initial baculovirus, subsequently amplified in a larger volume of Sf9 cells and used to infect at least 1 L of Sf9 cells harvested 72 hr post-infection. Cell pellets were resuspended in buffer (20 mM Tris pH 8.5, 200 mM NaCl, 4 mM DTT and 10 % (v/v) glycerol) supplemented with: cOmplete protease inhibitor (Roche), 1 mM PMSF, 0.5 % (v/v) tween-20. Cells were lysed through manual homogenisation and sonication. Resultant supernatant was clarified by centrifugation (4,000 rpm, 20 min) and applied to Strep-tactin resin (IBA). Recombinant protein was eluted by incubation with 3C protease. Resin was washed with anion exchange buffer (20 mM Tris pH 8.5, 4 mM DTT). Eluted protein was purified by anion exchange chromatography (BIRC6: monoQ, Cytiva; UBA6 and Apaf1: Resource Q, Cytiva) and further purified through size exclusion chromatography (BIRC6: Tricorn Superose 6, Cytiva; UBA6 and Apaf1: Superdex 200, Cytiva) in 20 mM HEPES pH 7.5, 200 mM NaCl, 4 mM DTT.

### Protein expression and purification from *E. coli*

Sequences were cloned into the following vectors. pOPINS (His_6_-SUMO): SMAC (a.a. 56-239, WT or monomeric mutant (V81D)), HtrA2 (a.a.134 - 458, S306A), XIAP (a.a.132-497), cIAP1 (a.a.177-618), CBL (a.a.49-435), BIRC7 (a.a.72-298), RNF4 (a.a.1-190), RNF144 (a.a.15-292), ARIH1 (a.a.95-400), NEDD4 (a.a.938-1319); pOPINB (His_6_-3C): Caspase-9 (single (D330A) and triple (E306A, D315A, D330A) mutants), Caspase-3 (WT or C136A), Caspase-7 (D206A or C186A), Caspase-8 (a.a. 217-479), UBE2Z, FAT10 (ϕιCys), UBE3C (aa 693–1083)); pOPINK (His_6_-GST-3C): CHIP (a.a. 21-303). Proteins were expressed in Rosetta (DE3) *E. coli* grown in 2xTY medium supplemented with 34 μg mL^-1^ chloramphenicol and 30 μg mL^-1^ kanamycin and induced with 400 μM IPTG at 18 °C overnight with addition of 100 µM Zn(SO_4_) where appropriate. Pellets were resuspended in lysis buffer (20 mM Tris pH 8.5, 300 mM NaCl, 2mM β-ME, 20 mM imidazole or, in the case of GST-CHIP, 20 mM Tris pH 8.5, 300 mM NaCl, 4 mM DTT) supplemented with: lysozyme, DNase, 1 µM pepstatin, 2 µM leupeptin and 1 mM PMSF, lysed by sonication followed by centrifugation (18,000 rpm, 30 min). Recombinant proteins were captured by immobilised metal affinity chromatography. Affinity tags were removed by incubation with SUMO protease or 3C protease overnight at 4°C and purified by anion exchange chromatography (Resource Q, Cytiva) eluting against a gradient of 1M NaCl. Proteins were further purified by size exclusion chromatography (Superdex 75 or 200, Cytiva) in 20 mM HEPES pH 7.5, 150 mM NaCl, 4 mM DTT. Ubiquitin, UBA1 and UBE2D2 were expressed and purified as previously described^42^

### Identification of BIRC6 interactors by mass spectrometry

HEK293F cells grown in FreeStyle^TM^ 293 Expression Media (Gibco), were washed with PBS and lysed in 25 mM HEPES pH 7.4, 100 mM NaCl, 1 mM DTT supplemented with 0.2 % (v/v) Igepal CA-630 (Sigma), 1 x cOmplete protease inhibitor tablet (Roche), 80 μM leupeptin and 40 μM pepstatin. Lysates were cleared by centrifugation (13,000 rpm, 15 min, 4°C) followed by ultracentrifugation (45,000 rpm, 30 min, 4°C). At least 7 mg of lysate protein was incubated for 1 hr at 4°C with Strep-tactin resin (IBA) uncoated or coated with 500 μg Strep-mScarlet-BIRC6. Following incubation, resin was washed and bound proteins eluted by resuspension in 2.5X Laemmli SDS sample buffer. Three biological repeats were performed. Eluted proteins were processed for mass spectrometry analysis using suspension traps (S-Traps). Proteins were reduced with 200 mM DTT in 0.1 M Tris pH 7.8, followed by alkylation with 200 mM iodoacetamide in 0.1 M Tris pH 7.8 in the dark. Samples were acidified by addition of 12 % phosphoric acid and captured on S-Trap^TM^ midi columns (C02-midi, ProtiFi). Columns were washed with 90 % methanol in 100 mM triethylammonium bicarbonate (TAEB) with centrifugation at 4000 g. Captured proteins were digested with trypsin (1:100 w/w) in 1 mM HCl overnight at RT. Peptides were eluted, dried and dissolved in Buffer A (98 % MilliQ-H_2_0, 2 % CH_3_CN and 0.1 % TFA). A non-saturating peptide amount (based on the intensity of the TIC chromatogram) of each biological repeat was injected for LC-MS/MS analysis on a Dionex Ultimate 3000 nano UPLC (Thermo Scientific) coupled to an Orbitrap Fusion Lumos Tribid mass spectrometer (Thermo Scientific). The instrument was operated in a data-independent acquisition (DIA) mode adapted from a previously described method^43^. Proteins were identified using the DIA-NN search engine (version 1.8), and label-free quantification was performed in Perseus (version 1.6.2.3). Volcano plots were generated using protein group intensities from DIA-NN and significant enrichment was plotted using a t-test with permutation FDR = 0.01 for multiple-test correction and s0 = 0.1 as cut-off parameters. The mass spectrometry proteomics data will be made available via the ProteomeXchange Consortium via the PRIDE partner repository^44^

### Preparation of fluorescently labelled proteins

Ubiquitin and FAT10 (ΔCys) both containing an N-terminal Cys and SMAC proteins with a C-terminal cysteine, were buffer exchanged into 20 mM HEPES pH 7.5, 200 mM NaCl. BDP-FL, Sulfo-Cyanine5 maleimide and Sulfo-Cyanine3 maleimide (Lumiprobe) and Alexa Fluor 555 (Invitrogen) were added in three times molar excess of the protein. Dye-protein mixture was incubated overnight at 4 °C and reaction was quenched by adding 1 mM β-ME. Remaining dye was separated from labelled protein by size exclusion chromatography (Superdex 75, Cytiva) equilibrated in 20 mM HEPES pH 7.5, 150 mM NaCl, 4mM DTT. Labelled SMAC N-terminal peptide (sequence AVPIAQKSEC(Cy3)-Am) was synthesised (Peptide Protein Research Ltd, UK).

### MicroScale Thermophoresis (MST) analysis

His_6_-tagged caspases were labelled using the His-tag labelling Kit RED-tris-NTA 2^nd^ generation (NanoTemper). Briefly, all His_6_-tagged caspases were measured to have affinities higher than 10 nM towards the dye. Therefore, 12.5 nM dye and 25 nM caspases were used as per manufacturer’s instructions. All MST measurements were conducted on a Monolith NT.115 (NanoTemper). Increasing concentrations of IAPs were titrated into 25 nM labelled caspases and incubated for 20 min, RT prior to measurement in MST/FP buffer: 20 mM HEPES pH 7.5, 150 mM NaCl, 5 mM DTT, 0.05 % (w/v) CHAPS and 0.04 mg mL^-1^ BSA. Two technical repeats were recorded with the MST power set to 40 %, and the laser turned on and off for 30 s and 25 s, respectively. Binding curves were fitted and affinities (Kd) were calculated using MO.Affinity Analysis software (NanoTemper).

### Fluorescence Polarisation (FP) analysis

FP assays were performed on a Hidex microplate reader using 535 (20) nm and 590 (10) nm excitation and emission channels respectively. Serial dilutions of BIRC6 were pipetted into a 384-well black low binding plate (Corning) containing a final concentration of 5 nM labelled-SMAC variants. For the competition assay, a dilution series of unlabelled SMAC (WT) was added to wells containing a final concentration of 3 nM labelled SMAC peptide and 30 nM BIRC6. All measurements were recorded in technical triplicate and are representative of at least two independent repeats. Data were plotted and fitted in GraphPad Prism 9.

### Ubiquitin ligase assays

Ubiquitin ligase assays were performed using 15 μM BDP-labelled ubiquitin, 0.3 µM E1 (UBA1 or UBA6), 0.9 µM E2 (UBE2D2 or BIRC6) and either 2 μM E3 (XIAP / cIAP1 / RNF4 / RNF144 / ARIH1 / NEDD4 / UBE3C) or 4 μM E3 (BIRC7 / GST-CHIP / CBL) under the following conditions: 40 mM HEPES pH 7.3, 10 mM MgCl_2_, 10 mM ATP, 0.6 mM DTT at RT. Samples were taken at designated time points, quenched in 4X LDS Sample buffer (Invitrogen) and analysed by SDS-PAGE and visualised by fluorescence (iBright, Invitrogen) and Coomassie staining.

### E1-E2 transthiolation assay

Assays testing transfer of activated ubiquitin from E1 ligases onto E2 ligases were performed using 0.5 µM E1 (UBA1 or UBA6), 3 µM E2 and 5 µM BDP-labelled ubiquitin or Cy3-labelled FAT10, in buffer conditions described in *Ubiquitin ligase assays*. Reactions were incubated for 30 min, RT and quenched and analysed as described in *Ubiquitin ligase assays*.

### Ubiquitination and FAT10ylation assays

Assays were performed in buffer conditions described in *Ubiquitin ligase assays* with 0.3 µM UBA6, 0.9 µM BIRC6 / XIAP / cIAP1, 15 μM BDP-labelled ubiquitin or Cy3-labelled FAT10 and 5-10 µM substrates. 0.9 µM UBE2D2 was included for XIAP and cIAP1 ubiquitination reactions. Samples were quenched and analysed as described in *Ubiquitin ligase assays*.

### Gel-based caspase-inhibition assay

Caspase activity assays were performed using caspase-9 variants at 1 µM (without apoptosome) and 0.5 μM (with apoptosome at a molar ratio of 1:5:10 caspase-9 : Apaf-1 : cytochrome C (SIGMA)) in the following buffer conditions: 25 mM HEPES pH 7.5, 100 mM KCl, 5 mM DTT. Caspase-9 variants were incubated for 10 min RT with IAPs at 6-fold molar excess before initiating the reaction by substrate addition (50 µM procaspase-3, C163A). Samples were quenched at time points as described in *Ubiquitin ligase assays* and analysed by SDS-PAGE and Coomassie staining.

### Caspase fluorogenic substrate cleavage assay

Fluorogenic cleavage assays to evaluate activity of caspases-3, −7 and −9 were performed in 384-well black low-binding plates (Corning) on a SpectraMax M3 (Molecular Devices) with excitation and emission wavelengths of 376 and 482 nm, respectively. Ac-DEVD-AFC (SIGMA) fluorogenic substrate was used for caspases-3 and −7 and Ac-LEHD-AFC (SIGMA) for caspase-9–apoptosome complex. Direct inhibition assays were performed with increasing concentrations of IAPs mixed with final concentrations of 5 nM caspase-3, 2.5 nM caspase-7 or 200 nM caspase-9–apoptosome complex in: 25 mM HEPES pH 7.5, 100 mM NaCl, 10 mM DTT, 0.01 % (v/v) Tween-20, and incubated for 20 min RT prior to measurement. Each reaction was mixed with 100 μM of the respective substrate. Initial rates were derived by calculating the gradient of the linear part of the curve. Analysis was performed in GraphPad Prism 9.4.1.

### Cryo-EM sample preparation and screening

3 µL BS3-cross-linked BIRC6-SMAC (1:3) or BIRC6 concentrated to 1.5 or 4 mg/mL was applied onto glow-discharged (20 mA, 30 sec) UltrAuFoil 1.2/1.3 Au 300 mesh grids and vitrified in liquid ethane using a Vitrobot Mark IV (Thermo Scientific), set to 4 °C and 100% humidity with a blotting force and time of −10 or 0 and 5 or 2.5 sec, respectively. BIRC6-SMAC grids were screened on a Talos Arctica (Thermo Scientific) with a Falcon 4 detector (Thermo Scientific).

### Data collection and processing of BIRC6

Cryo-EM data was collected on a Titan Krios G3 microscope (Thermo Scientific) equipped with a Gatan Quantum Image filter (20 eV slit width) and Gatan K3 direct electron detector. In total 4749 movies were collected at a pixel size of 0.43 Å/px in super-resolution counting mode using EPU’s (Thermo Scientific) faster-acquisition mode with a defocus range from −1 to −2.5 µm. Per foil hole two movies were acquired with a total dose of 49.9 e^-^/Å over a 3.5 sec exposure which were fractioned into 40 frames. Raw movies were clustered into optics groups using the extended file information (xml-files) of the EPU-data collection and the program EPU_TO_AFIS (https://github.com/DustinMorado/EPU_group_AFIS). Super-resolution movies were Fourier cropped 2x, motion corrected and dose-weighted with MotionCorr2 implementation in RELION3.1^45^. Contrast transfer function (CTF) of corrected micrographs were estimated with CTFFIND4.1^46^. Micrographs with a CTF estimation resolution worse than 6 Å were discarded for further processing. Particles were picked using the pre-trained ‘resnet8_u64’ and a self-trained (CNN: conv63, ∼1000 manually picked particles) TOPAZ model. Picked particles were extracted in 4x or 6x down-sampled 128 x 128 pixel boxes and classified with reference-free 2D classification in RELION3.1 for several rounds. Particles belonging to Class-averages showing high-resolution features were merged and duplicates were removed within a 100 Å cut-off radius. A total of 548,509 particles after 2D classification were used for initial 3D model generation in cryoSPARC v3.2^47^. All 548,509 particles and the initial model were used for 3D classification in RELION3.1. Particles (241,905) belonging to 3D-classes showing clear separation of all domains were selected, re-extracted in unbinned 640 x 640 pixel boxes and then auto-refined in RELION3.1 while applying C2 symmetry using a scaled, C2-aligned reference map from the output of the 3D Classification. Per-particle CTF parameters were optimised, and data-set wide optical aberrations were corrected through iterative rounds of CTF refinement^48^ followed by another round of 3D-Refinement. Further, Bayesian polishing^49^ with default parameters and an additional CTF refinement (including 4^th^ order aberration) yielded a map at 3.2 Å resolution using a soft mask according to the gold-standard criterion^50^. After another 3D classification in C2 symmetry without image alignment, 127,710 particles were chosen which yielded a reconstruction with a global resolution of 3.3 Å (EMD-15652). To improve the resolution of the flexible N terminus, specifically the DOC-like domain, and the C terminus, MultiBody refinement^51^ with masks covering 1) the central α-solenoid, 2) the N-terminal all-beta jelly-roll and DOC-like domains and 3) the C terminus was performed on the final symmetry expanded particle stack (255,420). MultiBody refinement on pre-CTF multiplied and symmetry expanded particles resulted in three new maps with resolutions of 2.9, 3.2 and 3.1 Å for the central core (EMD-15648), N terminus (EMD-15650) and C terminus (EMD-15651), respectively.

### Data collection and processing of BIRC6-SMAC

Data was collected on a Titan Krios G3 microscope (Thermo Scientific) with a Gatan Quantum Image filter (20 eV slit width) and Gatan K3 direct electron detector. In total 10,217 movies were collected at a pixel size of 0.829 Å/px in counting mode using the faster-acquisition mode of EPU (Thermo Scientific) with a defocus range from −0.75 to −2.5 µm. Per foil hole two movies were acquired with a total dose of 47.27 e^-^/Å over a 2.3 sec exposure which were fractioned into 50 frames. Movies were motion corrected and dose-weighted and CTFs of corrected micrographs were estimated with SIMPLE v3.0^52^. Next, template-based particle picking was used followed by two rounds of 2D classification in SIMPLE v3.0 and 1,210,317 particles were exported into cryoSPARC. Particles were classified into three classes during ab initio processing. A class containing 413,385 particles were used for non-uniform refinement in cryoSPARC^53^ using the volume generated in the previous ab initio stage. The particle stack and output poses of the non-uniform refinement were used in cryoDRGN^54^ version 0.3.4 for conformational heterogeneity analysis and subsequent classification of more distinct states. Particles were down sampled to a box size of 256 x 256 pixel (pizel size: 1.658 Å/px). A cryoDRGN model was trained using the highest architecture with three layers (1024×3) and an 8-dimensional latent variable Z for 25 epochs. To account for imperfect poses of the non-uniform refinement job we further included local pose refinement in cryoDRGN and 20 reconstructions were requested. Particles (36,872) corresponding to the two best out of twenty representative density maps were selected and converted to RELION format. Particles were imported and re-extracted in cryoSPARC with a 512-pixel box size and subjected to non-uniform refinement. The final reconstructed map was obtained by Bayesian polishing and subsequent non-uniform refinement in cryoSPARC, yielding a global resolution of 3.0 Å according to the gold standard criterion of 0.143 (EMD-15654). Local resolution was estimated using RELION.

### Model building and refinement of BIRC6-SMAC

Twenty AlphaFold2^55^ predictions of 1,400 aa overlapping BIRC6 fragments from the AlphaFold protein structure database (https://www.alphafold.ebi.ac.uk/)40,56 were separately rigid body fitted into the cryo-EM maps generated by MultiBody refinement through molrep (CCP4 suite)^57^. Correctly placed models were adapted to the respective map using Coot v0.9^58^. The structure was further refined by PHENIX real-space refinement against a composite map (EMD-15653) of all three MultiBody maps. Atomic models for SMAC homodimer (aa 56-239) and the N terminus of BIRC6 (aa 1-1700) were predicted using AlphaFold2^55^ via the ColabFold notebook^59^. The highest-ranking models were selected and, after removing disordered regions, rigidly docked into one of the two cryoDRGN reconstructions using UCSF ChimeraX^60^. The BIRC6 N termini and SMAC homodimer were then conformationally refined using Molecular Dynamics Flexible Fitting^61^ in NAMD v2.14^62^ and the resulting coordinates were combined with the model of the BIRC6 α-solenoid core. The entire structure was further refined using ISOLDE^63^ in ChimeraX, correcting highlighted stereochemical outliers where possible, followed by PHENIX real-space refinement^64^. Models were validated through MolProbity^65^ in PHENIX.

### Modelling and molecular dynamics simulation of BIRC6 BIR-SMAC AVPI sequence

An initial model of BIRC6 BIR bound to AVPI peptide was constructed by combining the AlphaFold2 prediction of BIRC6 BIR with peptide coordinates resulting from an overlay of the XIAP BIR2 AVPI-bound structure (PDB 4J46)^66^. A Zn^2+^ ion bound to XIAP BIR2 was also included in the BIRC6 BIR model and cysteine residues in the zinc binding site were deprotonated accordingly. The structure was then solvated with TIP3P water molecules and 150 mM NaCl and subjected to a series of conjugant gradient energy minimizations followed by two 5 ns equilibration simulations, first with harmonic restraints on the backbone atom positions and then on the alpha carbon positions. Finally, a 100 ns production simulation without restraints was carried out to ensure peptide stability. All molecular dynamics simulations were carried out using NAMD v2.14^62^ and the CHARMM36 force field^67^, and were conducted in the NPT ensemble with conditions maintained at 1 atm and 310 K using the Nosé–Hoover Langevin piston and Langevin thermostat, respectively. The r-RESPA integrator scheme was employed with an integration time step of 2 fs and SHAKE constraints applied to all hydrogen atoms. Short-range, nonbonded interactions were calculated every 2 fs with a cut-off of 12 Å; long-range electrostatics were evaluated every 6 fs using the particle-mesh-Ewald method.

## Acknowledgements

The authors would like to acknowledge A. von Kugelgen, L. Carrique, T. Matthews-Palmer, A. Costin and J. Caesar for help and advice for cryo-EM data collection and processing. Electron microscopy provision was provided through eBIC (proposal BI28731) and the Central Oxford Structural Molecular Imaging Centre (COSMIC) electron microscopy facility. Mass spectrometry analysis was performed in the Discovery Proteomics Facility (Target Discovery Institute) led by Roman Fischer / Iolanda Vendrell. We thank F. Barr, M. Higgins and S. Newstead for critical reading of the manuscript. L.D. and C.R. are funded through a Wellcome Trust studentship in Cellular and Structural Biology. A.P-F and B.M.K. are supported by the Chinese Academy of Medical Sciences (CAMS) Innovation Fund for Medical Science (CIFMS), China (grant number: 2018-I2M-2-002) and by Pfizer. P.R.E. is supported by a Medical Research Council (MRC) career development fellowship (MR/R008582/1).

## Author contributions

P.R.E designed and initiated the study. L.D performed cryo-EM sample preparation, data collection and analysis. L.D, C.J.E, C.R, D.F and P.R.E. performed experiments. Molecular dynamics simulations and flexible fitting were performed by K.C. A.P-F performed peptide in-solution digest and data analysis, supervised by B.K. P.R.E. wrote the manuscript.

## Data availability

All reagents are available under reasonable request from the corresponding author. Cryo-EM maps are deposited in the Electron Microscopy Data Bank under accession numbers: EMD-15648, 15650, 15651, 15652, 15653, 15654. Atomic coordinates were deposited in the PDB under accession number: 8ATM and 8ATO for BIRC6 and BIRC6–SMAC structures, respectively.

---

**Extended Data Fig. 1:**
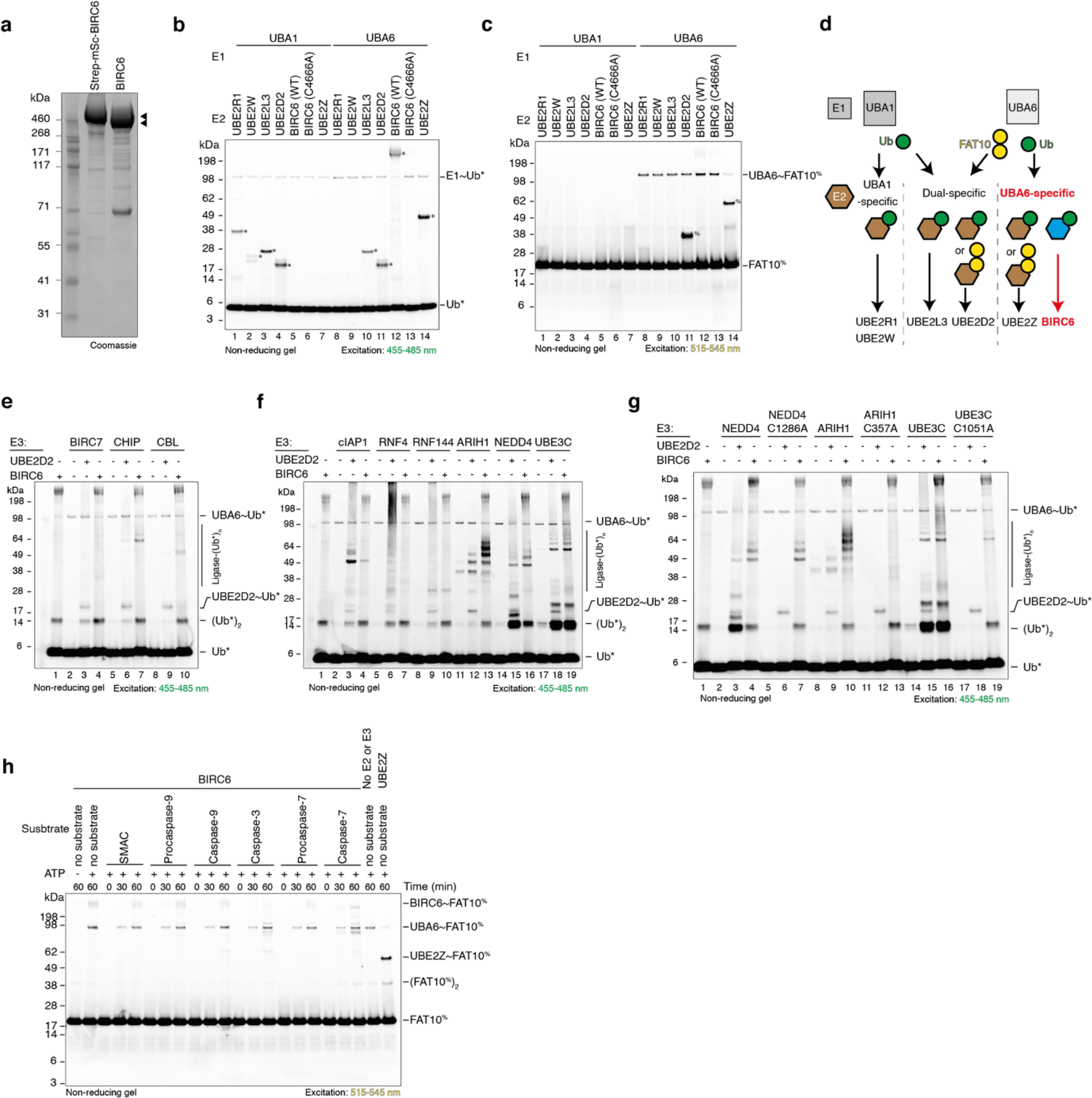
BIRC6 is a ubiquitin specific, cross-family E2 working exclusively with UBA6. **a**, SDS-PAGE gel analysis of full-length recombinant BIRC6 purified from Sf9 cells with a Twin-Strep-mScarlet tag followed by removal of the tag. **b - c**, BIRC6 accepts ubiquitin (**b, lane 12**) but weakly accepts FAT10 (**c, lane 12**) from non-canonical E1, UBA6, in an in vitro transthiolation reaction using BDP-labelled ubiquitin (*) or Cy3-labelled FAT10 (^%^) and quenched after 30 min. E2s known to only function with UBA1 (UBE2R1 and UBE2W) or both UBA1 and UBA6 (UBE2L3 and UBE2D2), were included for comparison. ∼ denotes a thioester linkage. **d**, Summary of ubiquitin/FAT10 transthiolation reactions between the two different E1s and a subset of E2s tested. Our results position BIRC6 as the only ubiquitin-specific E2 ligase working specifically with UBA6. **e**, BIRC6 ability to transfer ubiquitin onto a subset of the three different E3 ligase families were tested: BIRC7, cIAP1 and RNF4 - homodimeric Really Interesting New Gene (RING); CHIP - homodimeric U-Box RING; CBL - monomeric RING; RNF144 and ARIH1 – RING-in-between-RING (RBR); NEDD4 and UBE3C – Homologous to E6AP C-terminus (HECT). All reactions included UBA6 with either UBE2D2 or BIRC6 as the E2. **f**, BIRC6 acts as an E2 ligase and facilitates transfer of ubiquitin onto ARIH1 (**lane 13**), NEDD4 (**lane 16**) and UBE3C (**lane 19**) in auto-ubiquitination reactions with BDP-labelled ubiquitin (*). **g**, To distinguish ubiquitin transfer from ubiquitination of an E3 acting as a BIRC6 substrate, ubiquitin transfer to catalytically inactive NEDD4, ARIH1 and UBE3C was tested. BIRC6 catalyses ubiquitin transfer to ARIH1 (compare lanes 10 and 13) and UBE3C (compare lanes 16 and 19) but uses NEDD4 as a substrate (compare lanes 4 and 7). All reactions included UBA6 with either UBE2D2 or BIRC6 as the E2. **h**, BIRC6 does not transfer FAT10 onto substrates, tested by an in vitro FAT10ylation reaction using UBA6 and Cy3-labelled FAT10 (^%^). UBA6 alone or in the presence of UBE2Z are included as assay controls. Representative gels of at least 2 independent repeats are shown.

**Extended Data Fig. 2:**
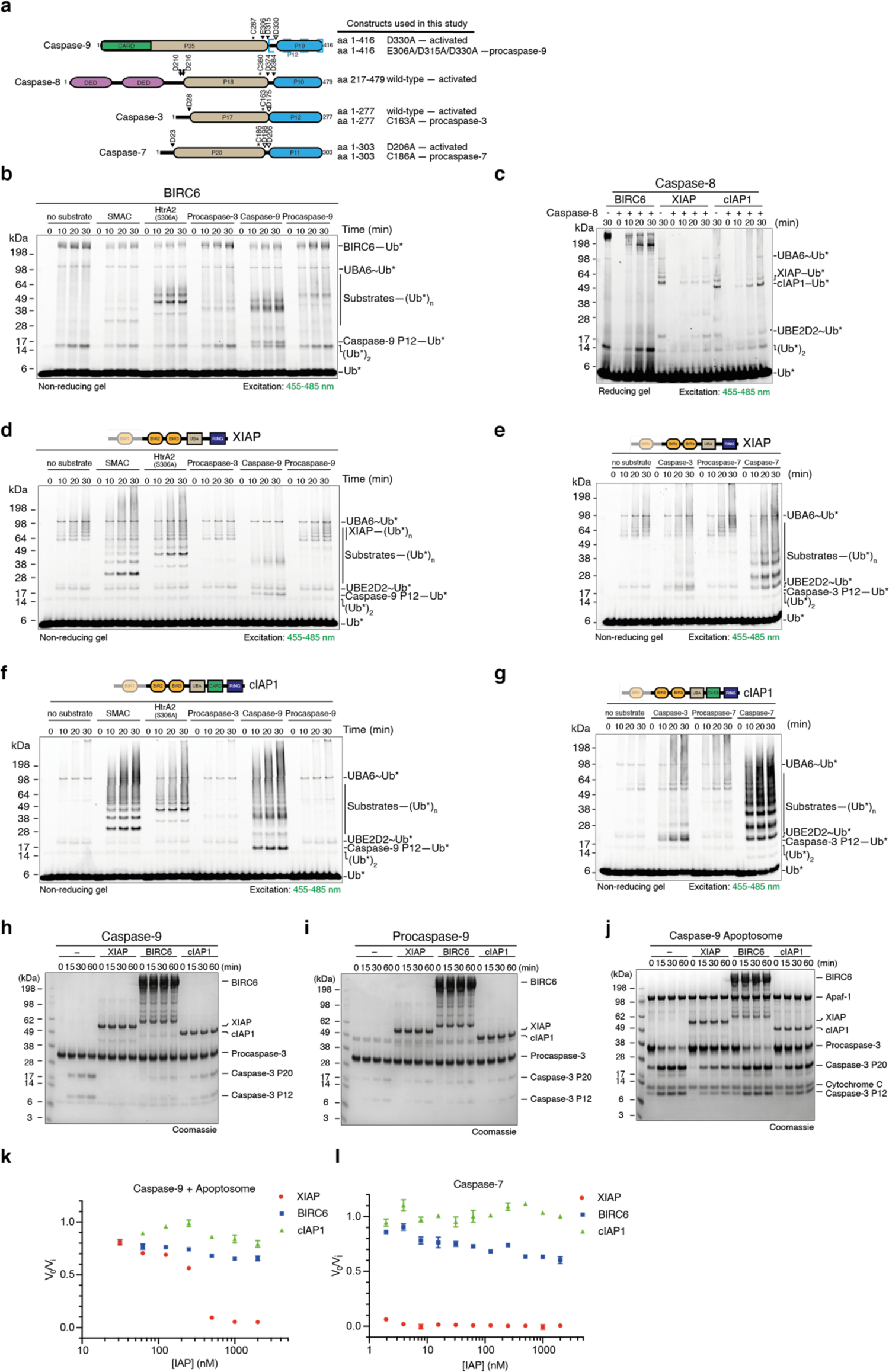
Comparison of ubiquitination and caspase inhibition activities of BIRC6, XIAP and cIAP1. **a**, Schematic depiction of caspase constructs used in our study. *denotes the catalytic cysteine; filled triangles represent autocleavage sites; empty triangles represent sites of cleavage by another caspase. Dashed blue box indicates caspase-9 P12 subunit. **b**, In vitro ubiquitination reaction testing BIRC6 ability to ubiquitinate intrinsic apoptosis regulators, SMAC and catalytically inactive HtrA2 (S306A), and caspases in the presence of UBA6 and BDP-labelled ubiquitin (*). **c**, In vitro ubiquitination reaction comparing activated caspase-8 ubiquitination by BIRC6, XIAP and cIAP1 in the presence of UBA6 and BDP-labelled ubiquitin (*). XIAP and cIAP1 reactions included the E2 UBE2D2. **d-g**, In vitro ubiquitination activity of XIAP and cIAP1 in the presence of UBA6, UBE2D2 and BDP-labelled ubiquitin (*) against **(d, f)** SMAC, HtrA2, activated caspases-3 and −9 and procaspase-9 or **(e,g)** activated caspases-3 and −7 and procaspase-7. Gels are representative of two independent repeats. **h-j**, Ability of activated caspase-9, procaspase-9 and activated caspase-9-apoptosome complex to cleave procaspase-3 in the absence or presence of XIAP, BIRC6 or cIAP1 in 6-fold excess. Gel images are representative of 5 independent repeats. **k & l**, BIRC6 weakly inhibits activated caspase-9 in the presence of the apoptosome complex, **k**, and activated caspase-7 activity, **l**, as measured in a fluorogenic substrate cleavage assay. Graph shows the rate of substrate cleavage by caspases in presence of IAP normalised against caspase activity alone. Data shows 2 independent triplicate repeats. Error bars represent SD.

**Extended Data Fig. 3:**
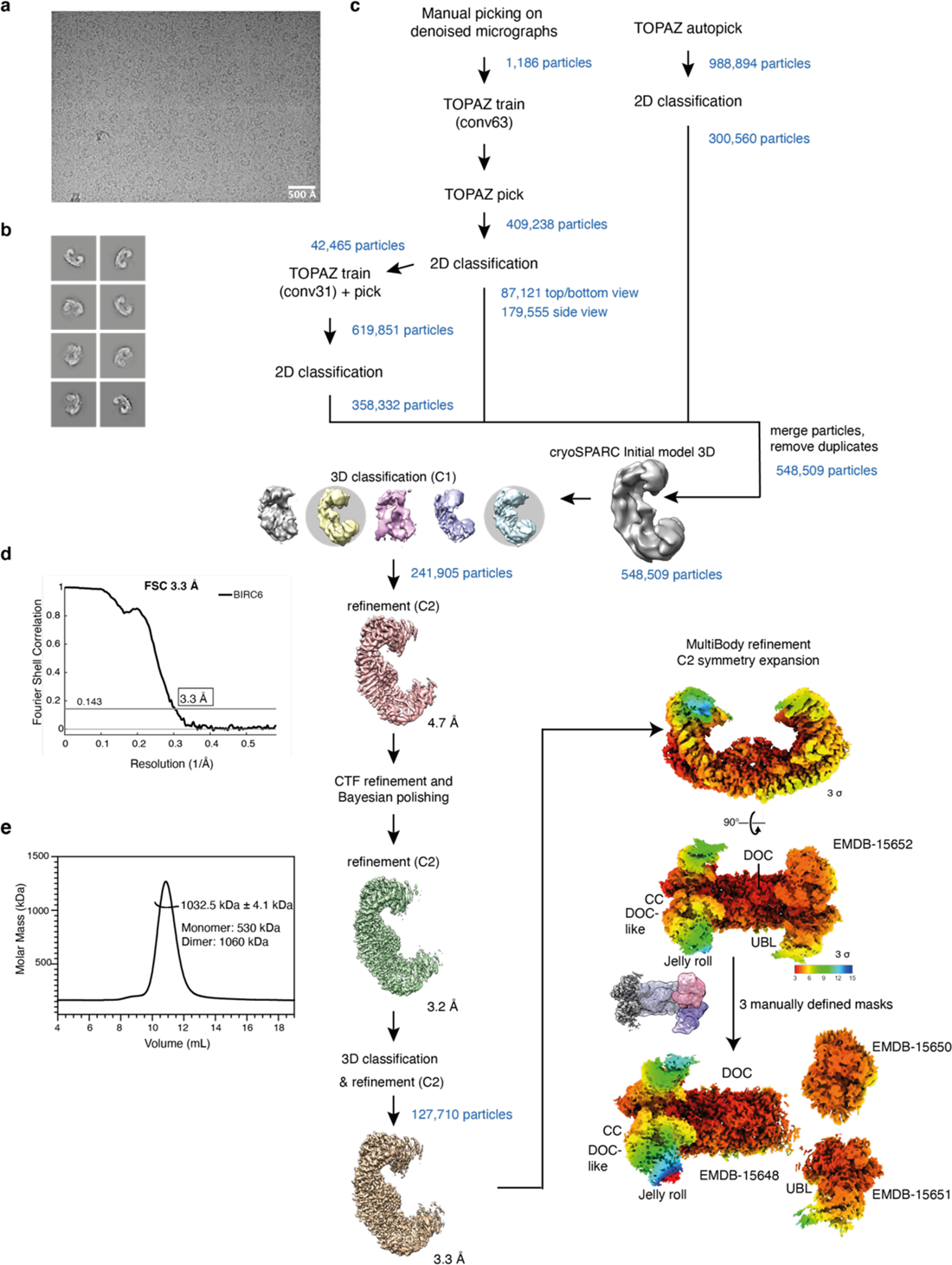
Cryo-EM processing workflow of BIRC6 structure. **a**, Representative micrograph of BIRC6 on UltrAuFoil R1.2/1.3 mesh 300 grids at 4 mg mL^-1^, pixel size: 0.68 Å. **b**, Representative 2D classes of BIRC6, box size: 250 Å. **c**, Image processing workflow for BIRC6. The final reconstruction of a consensus refinement (EMD-15652) yielded a global resolution of 3.3 Å. Local resolution of the consensus refinement cryo-EM map of BIRC6, global resolution: 3.3 Å. Subsequent MultiBody refinement with a C2 symmetry expansion applied and three manually defined masks, improved the density for the outward facing N and C termini (EMD-15650 and 15651, respectively). **d**, Fourier Shell Correlation (FSC) curve of the consensus refinement showing a global resolution of 3.3 Å. **e**, SEC-MALS of BIRC6 on a Superose6 column confirming dimerization of BIRC6 in solution, rendering it a 1 MDa complex.

**Extended Data Fig. 4:**
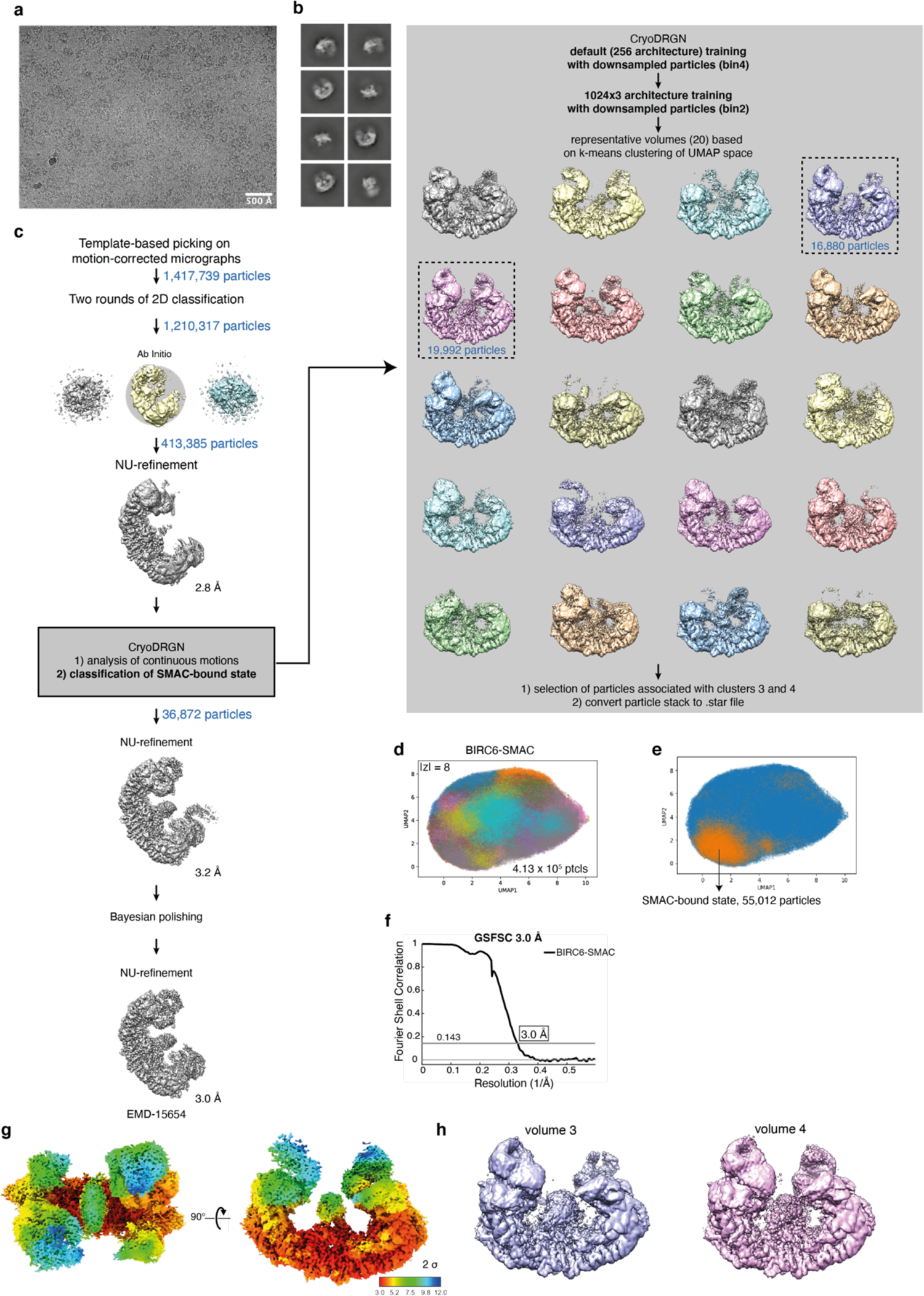
Cryo-EM processing workflow of BIRC6-SMAC structure. **a**, Representative micrograph of BIRC6 in complex with SMAC on UltrAuFoil R1.2/1.3 mesh 300 grids at 1.5 mg mL^-1^, pixel size: 0.68 Å. **b**, Representative 2D classes of BIRC6 in complex with SMAC, box size: 250 Å. **c**, Image processing workflow for BIRC6-SMAC structure. Whilst a reconstruction of cryoSPARC non-uniform refinement map yielded a global resolution of 2.8 Å, density for SMAC was only detected at lower resolutions. CryoDRGN was used to re-classify particles into classes. Classes three and four, containing the central SMAC density, were selected. The final reconstruction from non-uniform refinement yielded an overall resolution of 3.0 Å (EMD-15654). **d**, UMAP-representation of 413,000 BIRC6-SMAC particles after cryoDRGN training. Colours indicate 20 clusters based on k-means clustering. **e**, UMAP-representation of 55,012 BIRC6-SMAC particles (orange) containing SMAC bound to BIRC6 from classes 3 and 4 of the k-means clustering. **f**, Fourier Shell Correlation (FSC) curve of the consensus refinement showing a global resolution of 3.0 Å. **g**, Local resolution of the consensus refinement cryo-EM density of BIRC6, global resolution: 3.0 Å. **h**, cryoDRGN density reconstruction of classes 3 and 4 of the k-means clustering showing SMAC bound in the central cavity of BIRC6.

**Extended Data Fig. 5:**
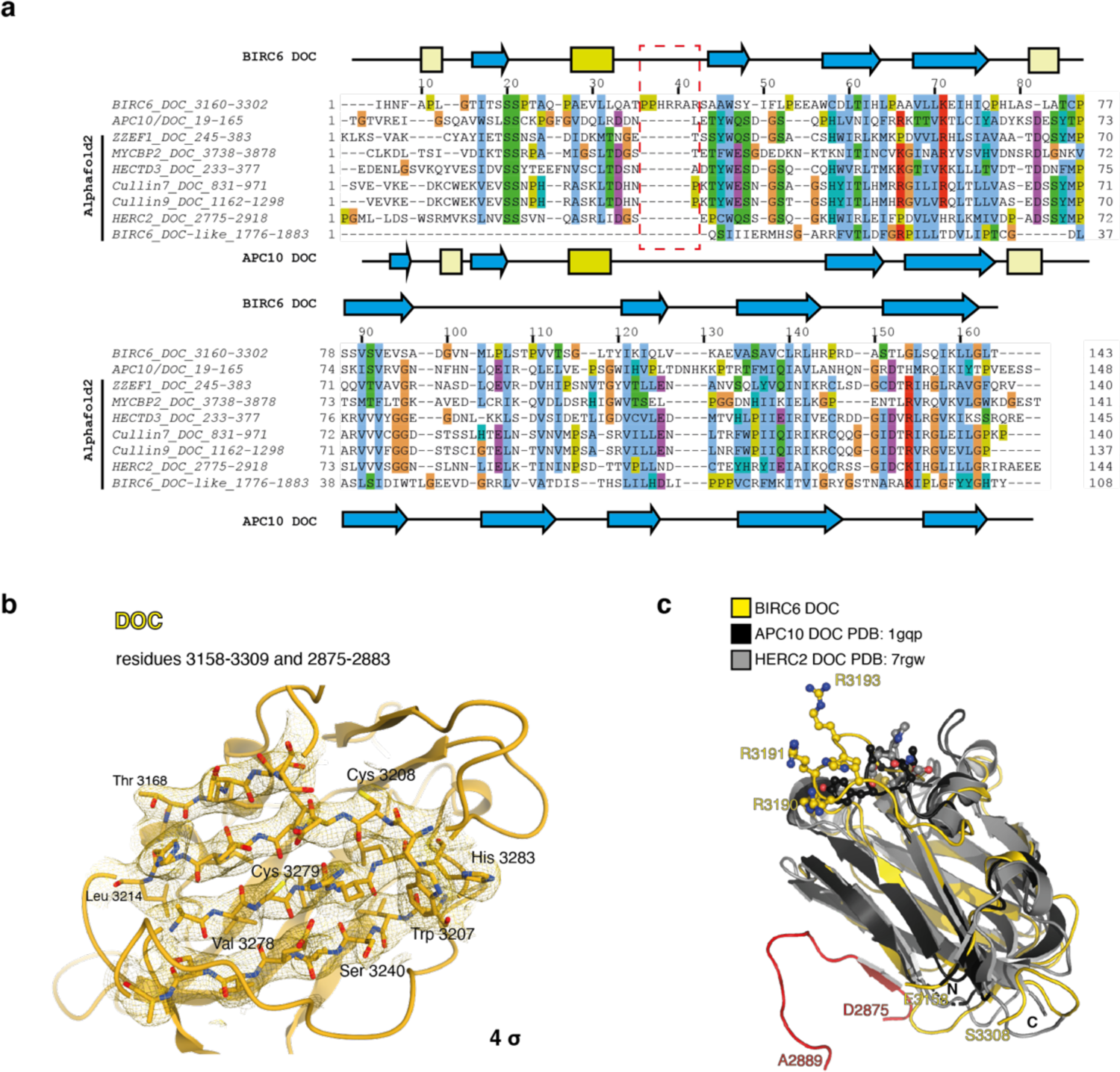
BIRC6 DOC domain includes a unique loop insertion with a positively charged tip. **a**, Structure-based sequence alignment coloured by residue chemistry of nine DOC domains with the secondary structure elements of BIRC6 DOC and APC10 DOC annotated above and below the sequences, respectively. Red box emphasises the loop insertion unique to the BIRC6 DOC domain. Alphafold2 was used to model DOC domains for which no structures are currently available. **b**, Representative cryo-EM density of the central DOC domain at 4 σ. A subset of well-defined residues is highlighted. **c**, BIRC6 DOC domain (yellow) superimposed onto DOC domains of APC10 (black, PDB: 1GQP) and HERC2 (grey, PDB: 7RGW). Red denotes residues Asp2875 – Ala2889 rising from the BIRC6 α-solenoid core to stabilise the DOC domain. Positively charged residues at the tip of the BIRC6 DOC domain loop are shown in atom representation.

**Extended Data Fig. 6:**
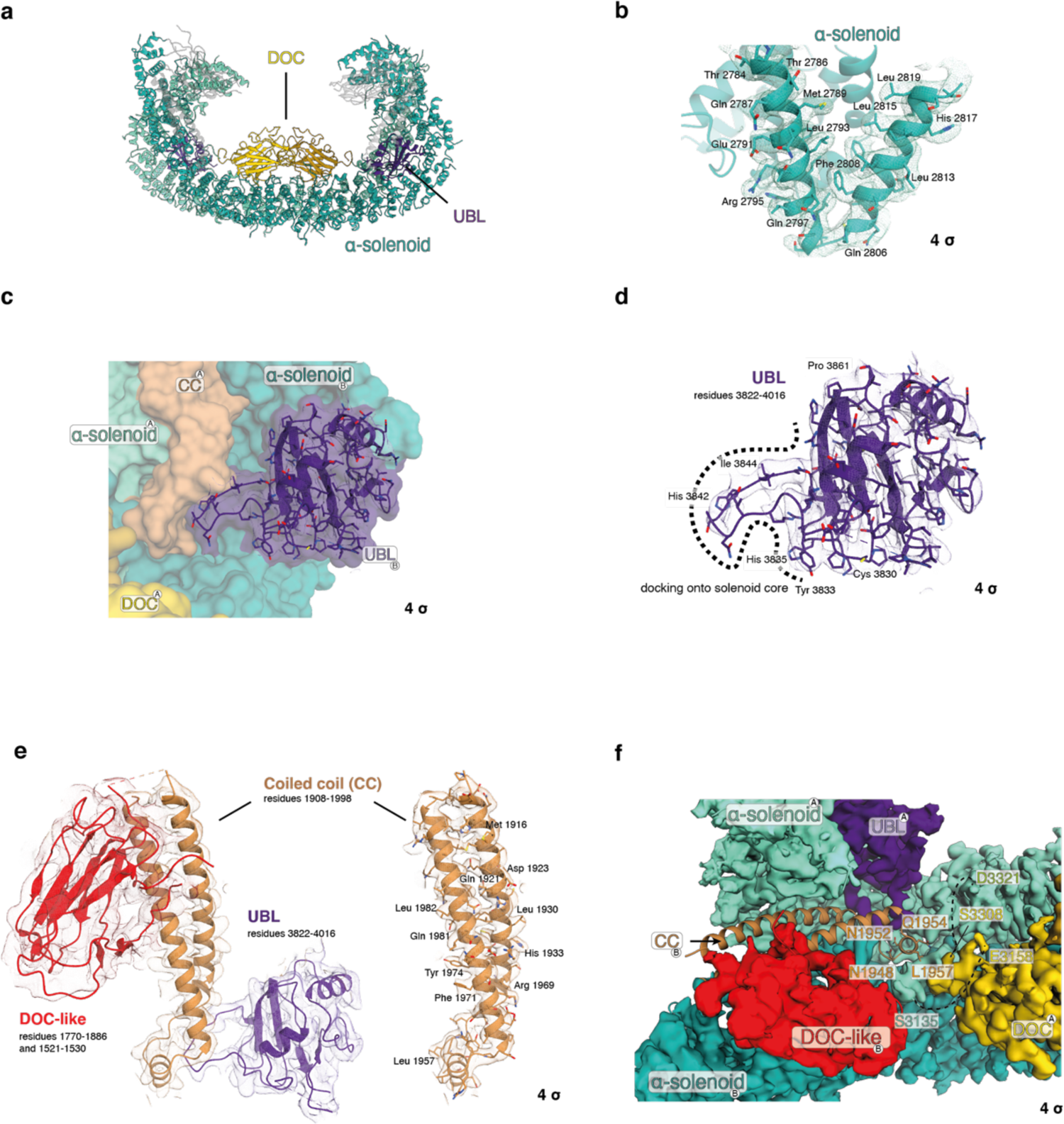
Representative cryo-EM density maps of BIRC6. **a**, Side view of BIRC6 in cartoon representation with the α-solenoid (aquamarine), DOC (gold) and UBL (purple) domains highlighted. **b**, Representative cryo-EM density at indicated σ levels in the α-solenoid core. **c**, Positioning of UBL domain in relation to α-solenoid core and coiled-coil (bronze) domain. Representative cryo-EM density at indicated σ levels in **d**, UBL domain, **e**, DOC-like (red), coiled coil and UBL domains. **f**, Coiled-coil domain acts as a structural hinge nestled between the α-solenoid core and interacting with the DOC-like and UBL domains. Chains to which the domains belong are indicated by ^A^ and ^B^.

**Extended Data Fig. 7:**
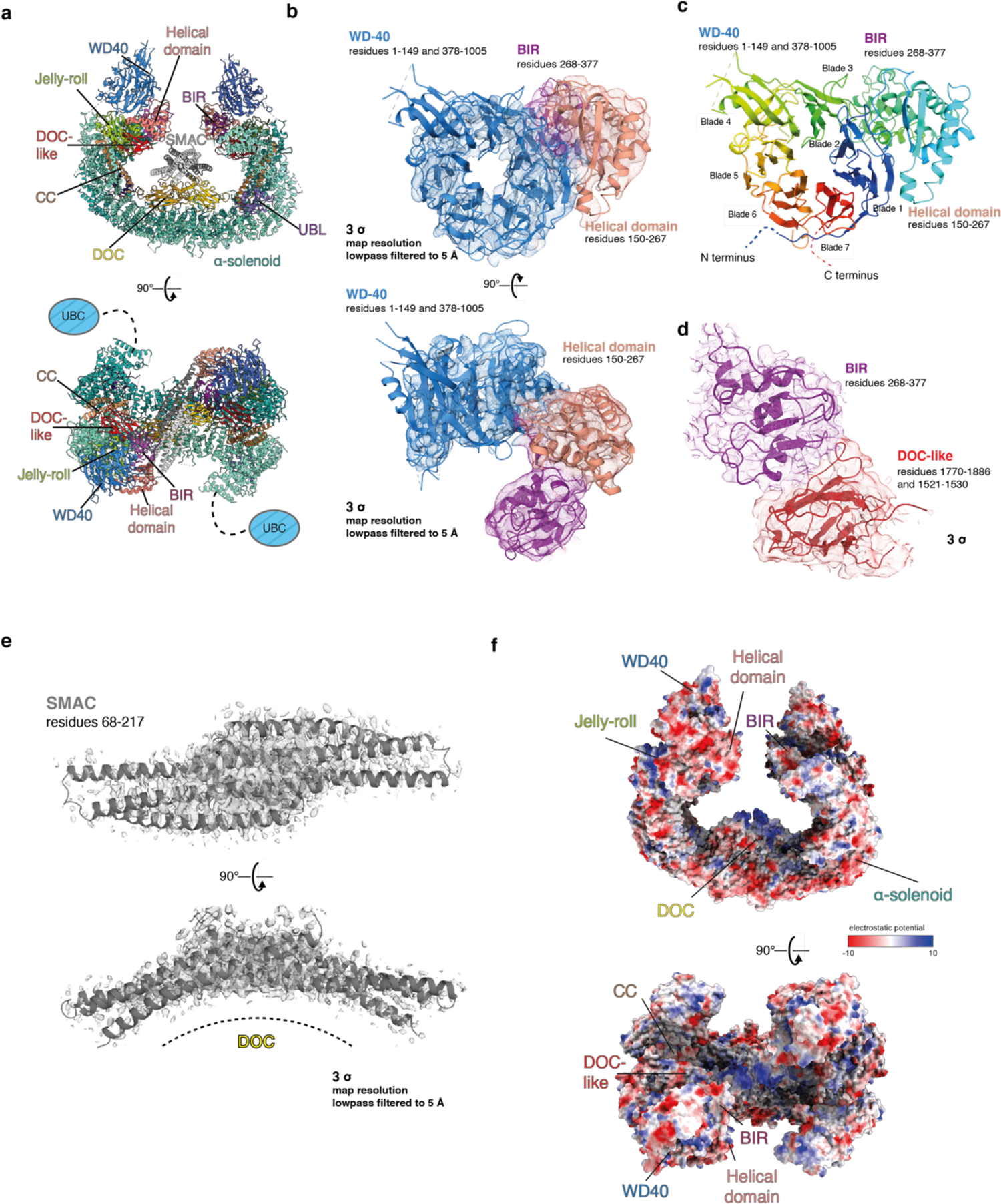
Representative cryo-EM density maps of BIRC6-SMAC complex. **a**, BIRC6 in complex with SMAC shown by cartoon representation. **b**, Representative cryo-EM density at indicated σ level in WD40, BIR and helical domain. **c**, Rainbow representation of the WD40, BIR and helical domains showcasing the BIR and helical domain insertions between the seven WD40 blades. N and C termini are indicated by blue and red dashed lines, respectively. Representative cryo-EM density at indicated σ levels in **d**, BIR and DOC-like domains and **f**, SMAC dimer. **e**, Electrostatic potential of BIRC6 shown as side and top orientation. Negatively and positively charged residues are shown in red and blue, respectively.

**Extended Data Figs. 8:**
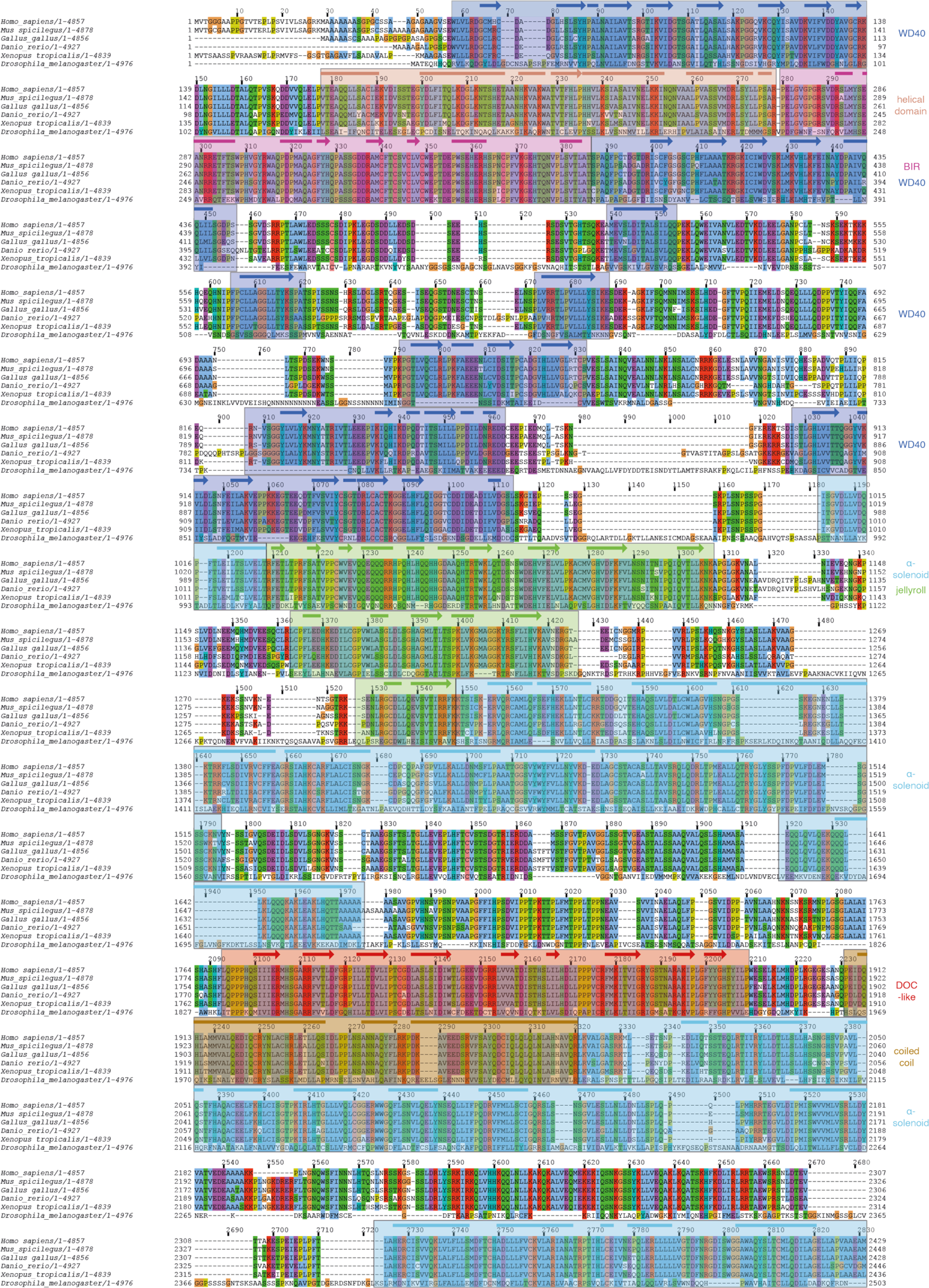
Structural features identified are conserved in BIRC6 orthologues Full-sequence alignment of six representative orthologues of BIRC6 coloured by residue chemistry using Jalview. *Mus spicilegus* (steppe mouse); *Gallus gallus* (Red junglefowl - chicken); *Danio rerio* (Zebrafish); *Xenopus tropicalis* (Western clawed frog); and *Drosophila melanogaster* (common fruit fly) are shown. Sequence regions corresponding to where cryo-EM density allowed for model building are highlighted in their respective domain colour. Arrows and bars above the sequence represent α-helical and β-sheet secondary structure elements, respectively. Domains are specified on the right.

**Extended Data Figs. 9:**
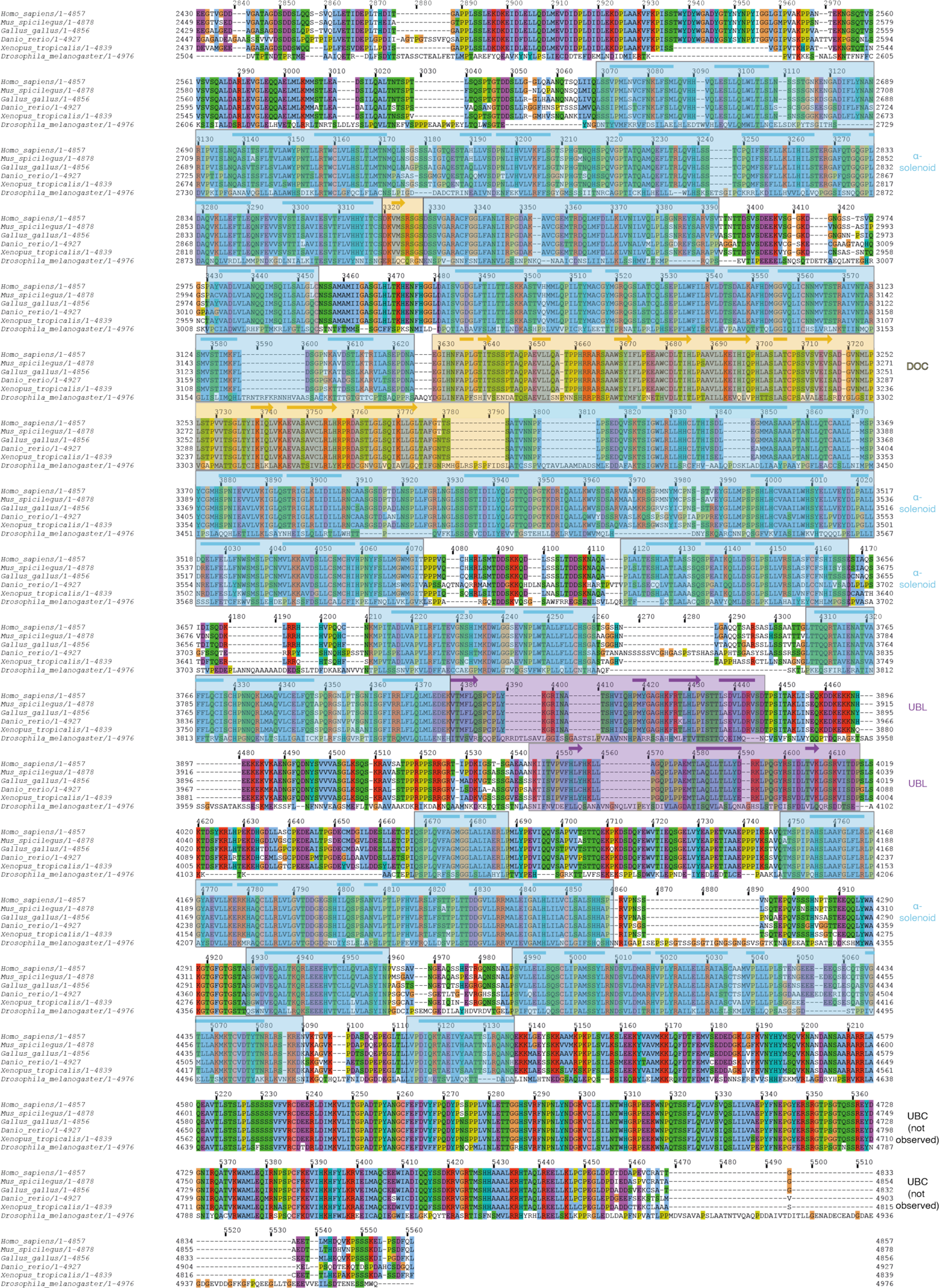
Structural features identified are conserved in BIRC6 orthologues Full-sequence alignment of six representative orthologues of BIRC6 coloured by residue chemistry using Jalview. *Mus spicilegus* (steppe mouse); *Gallus gallus* (Red junglefowl - chicken); *Danio rerio* (Zebrafish); *Xenopus tropicalis* (Western clawed frog); and *Drosophila melanogaster* (common fruit fly) are shown. Sequence regions corresponding to where cryo-EM density allowed for model building are highlighted in their respective domain colour. Arrows and bars above the sequence represent α-helical and β-sheet secondary structure elements, respectively. Domains are specified on the right.

**Extended Data Fig.10:**
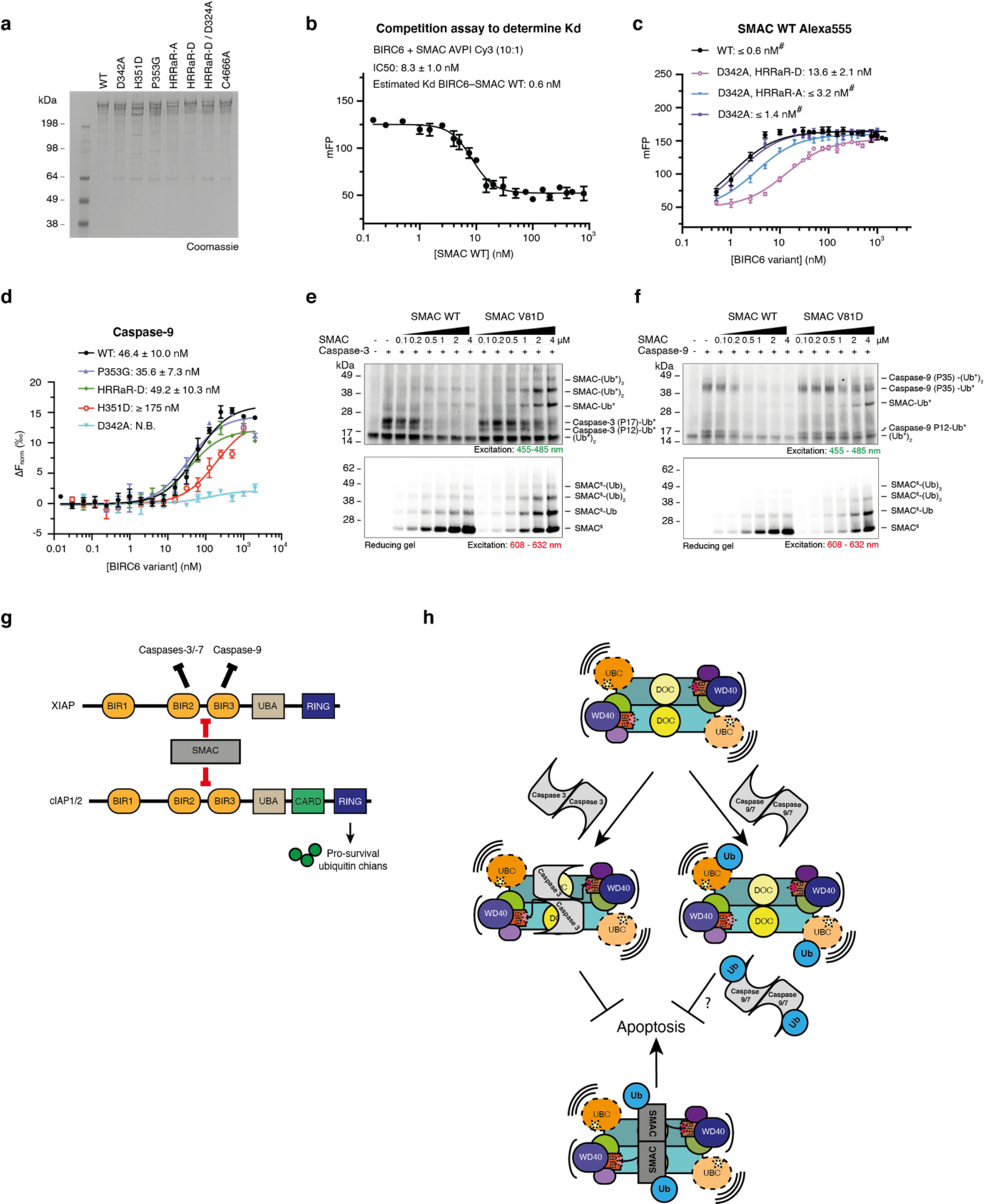
Dimeric SMAC antagonises BIRC6 through a sub-nanomolar affinity interaction out competing caspases. **a**, SDS-PAGE gel showing purified BIRC6 mutants used in this study. **b**, Estimation of sub-nanomolar affinity between BIRC6 and dimeric SMAC (wild-type, WT) using an FP competition assay titrating increasing concentrations of unlabelled WT SMAC against 30 nM BIRC6 and 3 nM Cy3-SMAC peptide. Data represents 2 independent repeats in triplicate. Affinity of WT SMAC and BIRC6 was estimated as 0.6 nM using an IC50 of 8.3 nM fitted to a tight binding equation^68^. **c**, FP assays determining affinities between BIRC6 WT or variants and Alexa555-SMAC (WT). # denotes the BIRC6-SMAC (WT) affinity being tighter than the minimum concentration of labelled SMAC detectable by FP, preventing accurate measurement of this affinity. Results are representative of 3 independent repeats in triplicate. **d**, Affinities between BIRC6 (WT) or variants and RED-tris-NTA labelled activated His_6_-caspase-9 measured using MST. Results are representative of 2 independent repeats in triplicate. Error bars represent SD. **e,f**, In vitro ubiquitination assays testing effect of increasing SMAC (WT) or SMAC monomer (V81D) concentration on BIRC6 ubiquitination of activated **e**, caspase-3, **f**, caspase-9 using BDP-labelled ubiquitin (*) and Cy-5-labelled SMAC variants (^$^). Assay quenched after 30 min. Gels are representative of 2 replicates and correspond to **Fig. 4d**. **g**, Summary of characterised anti-apoptotic activity of IAP members XIAP and cIAP1/2. XIAP directly inhibits caspases-9,-3 and −7 through binding via specific BIR domains. cIAP1/2 inhibits apoptosis indirectly by assembling pro-survival ubiquitin chains via its RING domain. Both IAP members are regulated by SMAC. **h**, BIRC6 comprises an anti-parallel dimer with crab-like architecture in which the flexible N-terminal substrate-binding modules acts as antennae recruiting substrates, such as activated caspases, positioning them on the highly conserved central DOC domain in the mouth-like cavity of BIRC6 for ubiquitination by highly flexible C-terminal catalytic claws. BIRC6 directly inhibits caspase-3 activity and multi-monoubiquitinates active caspases potentially priming ubiquitin chain extension by other E3 ligases leading to caspase degradation. SMAC competitively displaces caspases to disrupt BIRC6-caspase inhibition thereby antagonising BIRC6 anti-apoptotic activity.

**Supplementary File 1. Table 1:**
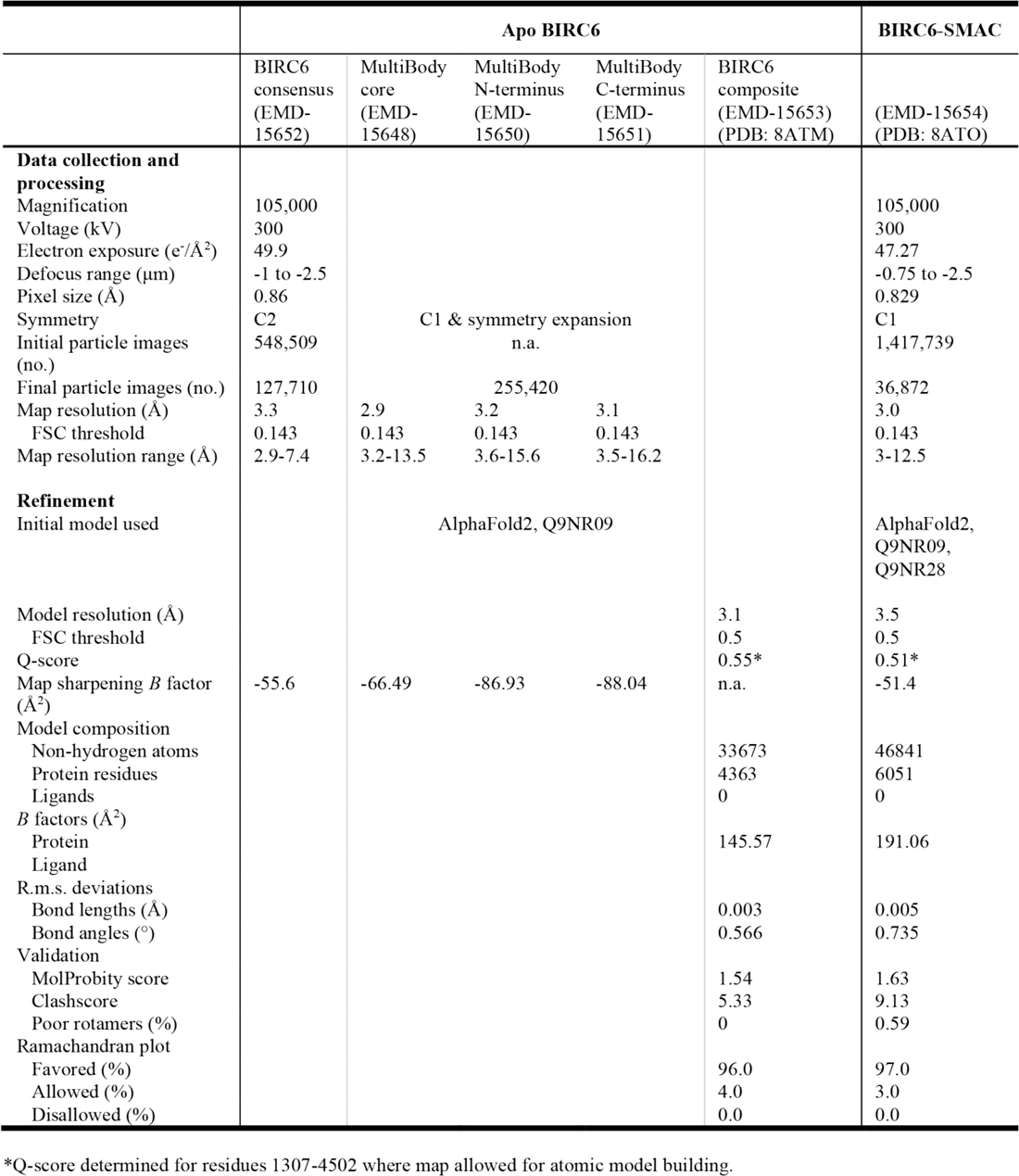
Cryo-EM data collection, refinement and validation statistics

**Supplementary file 2.**
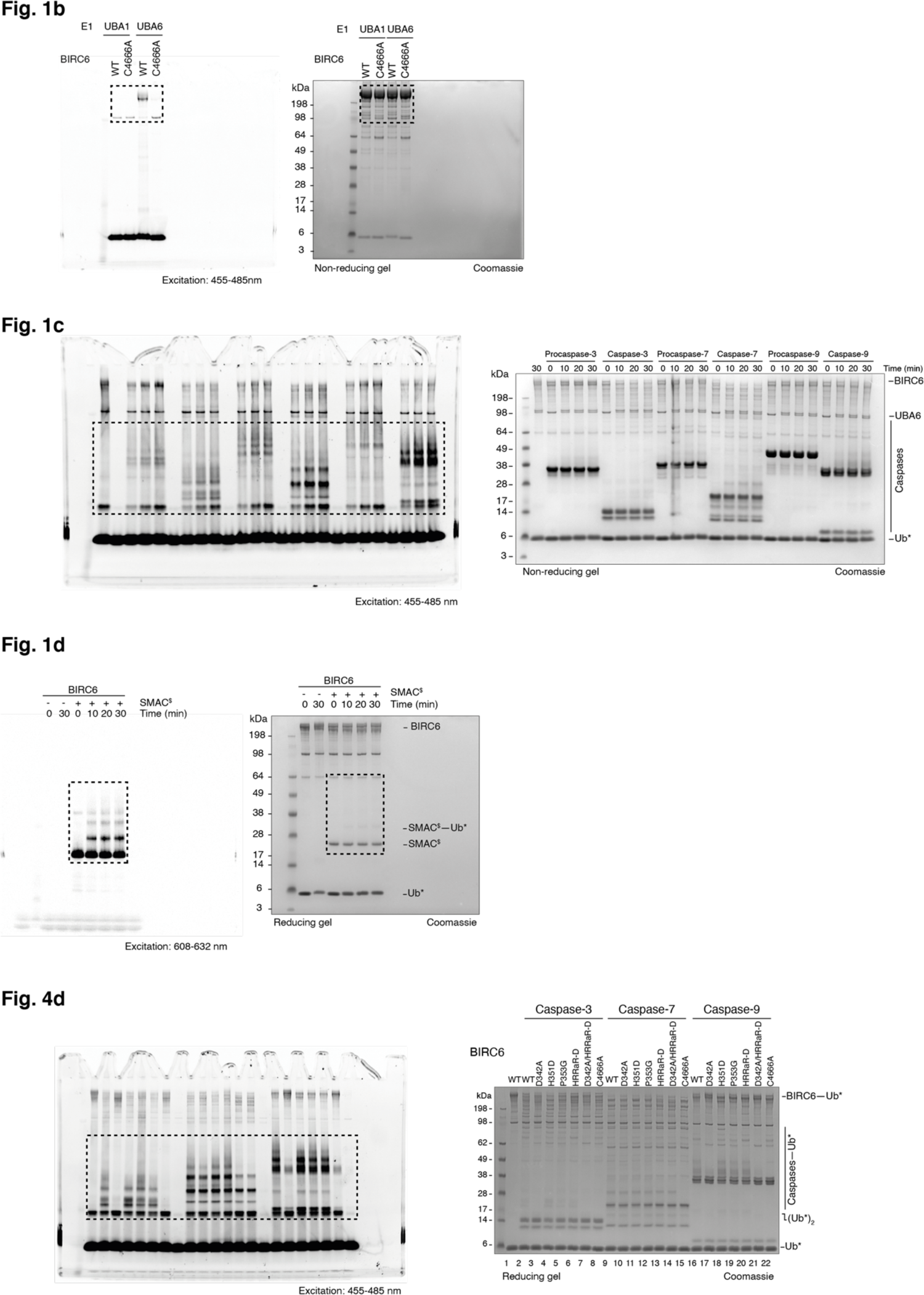
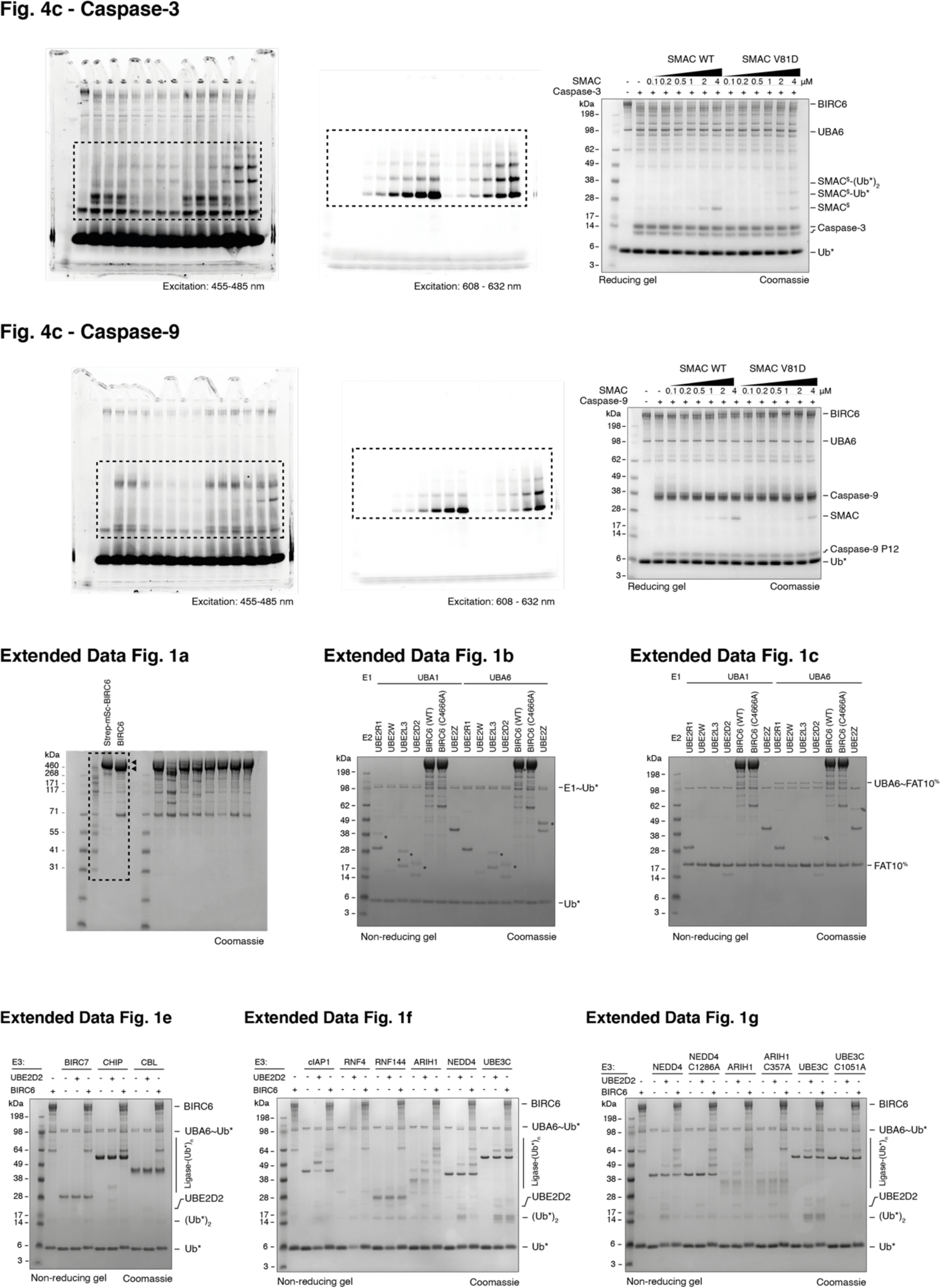
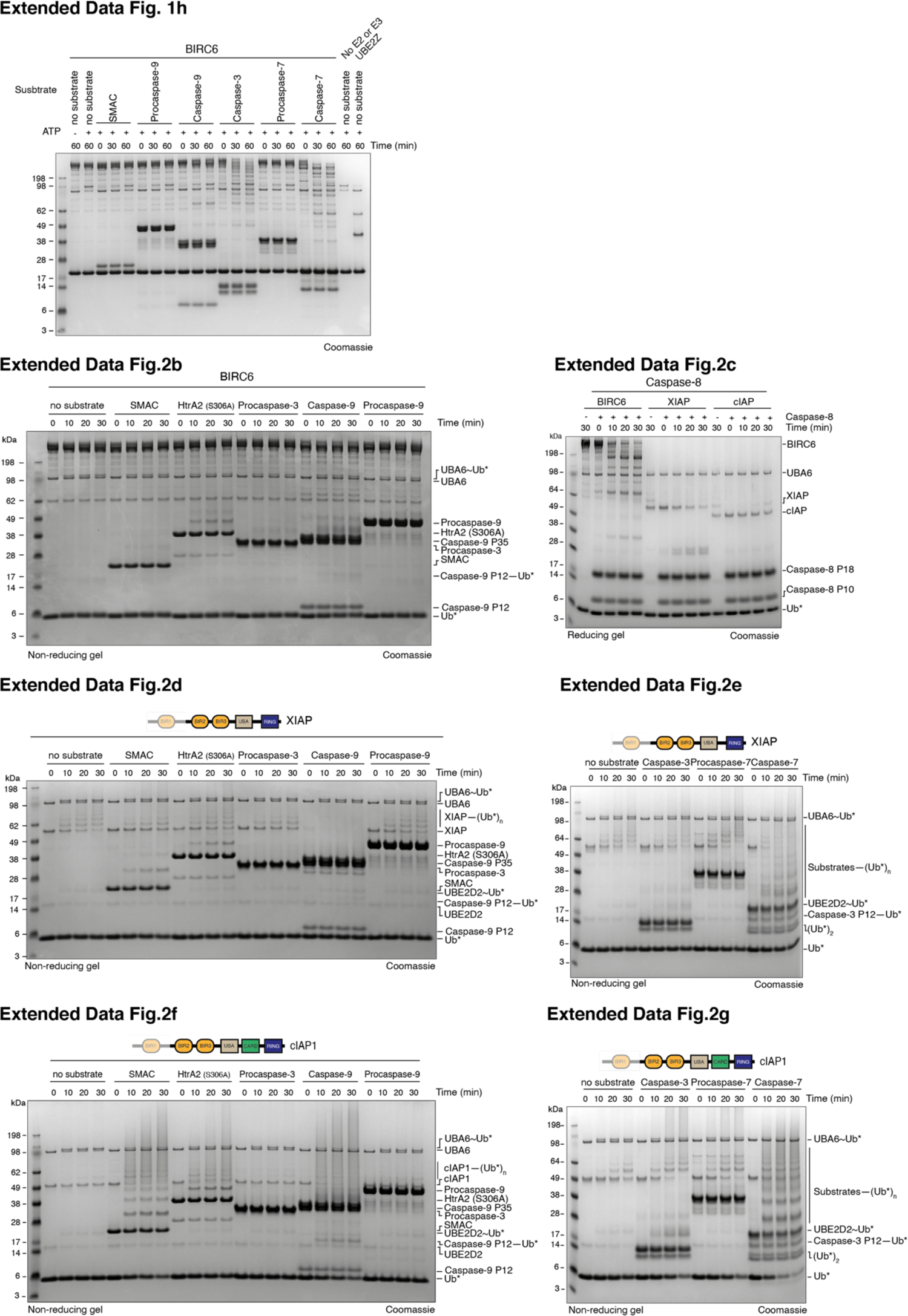
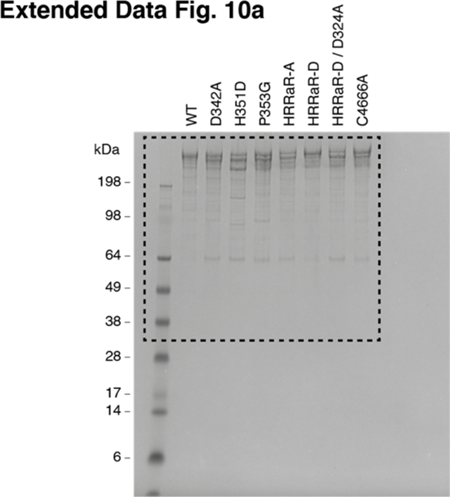
Uncropped fluorescence images and corresponding Coomassie stained gels

